# Broad host tropism of ACE2-using MERS-related coronaviruses and determinants restricting viral recognition

**DOI:** 10.1101/2022.09.11.507506

**Authors:** Chengbao Ma, Chen Liu, Qing Xiong, Mengxue Gu, Lulu Shi, Chunli Wang, Junyu Si, Fei Tong, Peng Liu, Meiling Huang, Huan Yan

**Author notes:** These authors contributed equally (Chengbao Ma, Chen Liu, Qing Xiong).

## Abstract

Phylogenetically distant coronaviruses have evolved to use ACE2 as their common receptors, including NL63 and many Severe acute respiratory syndrome (SARS) coronavirus-related viruses. We recently reported two Middle East respiratory syndrome coronavirus (MERS-CoV) closely related bat merbecoviruses, NeoCoV and PDF-2180, use Angiotensin-converting enzyme 2 (ACE2) for entry. However, their host range and cross-species transmissibility remain unknown. Here, we characterized their species-specific receptor preference by testing ACE2 orthologs from 49 bats and 53 non-bat mammals. Both viruses exhibited broad receptor recognition spectra and are unable to use ACE2 orthologs from 24 species, mainly Yinpterochiropteran bats. Comparative analyses of bat ACE2 orthologs underscored four crucial host range determinants, all confirmed by subsequent functional assays in human and bat cells. Among them, residue 305, participating in a critical interaction, plays a crucial role in host tropism determination. NeoCoV-T510F, a mutation that enhances human ACE2 recognition, further expanded the potential host range via tighter interaction with an evolutionary conserved hydrophobic pocket. Our results elucidated the molecular basis for the species-specific ACE2 usage of MERS-related viruses across mammals and shed light on their zoonotic risks.

## Introduction

The coronaviruses associated with human emergence in the past two decades impose severe threats to human health, especially the recent COVID-19 ^1–4^ As important coronavirus reservoirs, bats have been reported natural hosts of ancestors and relatives of three high-risk human β-CoVs: SARS-CoV, SARS-CoV-2, and MERS-CoV ^1,2,5–10^. The MERS-CoV belongs to the lineage C of β-CoVs (*Merbecovirus* subgenus) with a high case-fatality rate of 34.5%, according to the most recent update of the MERS situation of the World Health Organization (WHO) ^11^. NeoCoV and PDF-2180 are MERS-related viruses sampled in vesper bats harbor in South Africa and Southwest Uganda, respectively ^12,13^. NeoCoV represents the yet-identified closest relative of MERS-CoV with ~85% whole genome nucleotide similarity ^14^ However, the receptor binding domains (RBD) in spike protein (S) of the two viruses are very different from MERS-CoV and many other merbecoviruses, indicative of unique receptor usage ^12,15^.

ACE2 mediates viral entry of many SARS-related CoVs (*Sarbecovirus* subgenus), such as SARS-CoV, SARS-CoV-2, and bat coronavirus RaTG13 ^2,16,17^ Moreover, the α-CoV NL63 and their bat relatives also engage ACE2 for entry ^18^. In both cases, the viruses bind to a similar surface of the ACE2 protease domain, albeit through two groups of structurally distinct RBDs ^19^. Several merbecoviruses use their host’s DPP4 as entry receptors, such as MERS-CoV, bat CoV HKU4, and HKU25, whereas the receptors for many other merbecoviruses remain elusive ^12,20–22^. Recently, we reported that NeoCoV and PDF-2180 unexpectedly engage bat ACE2 as their receptors ^15^. Cryo-electron microscopy (Cryo-EM) analysis of NeoCoV or PDF-2180 RBD in complex with a bat ACE2 from *Pipistrellus pipistrellus* (Ppip) revealed a relatively small ACE2 binding surface featured by an N-glycosylation mediated protein-glycan interaction, a mode distinct from other ACE2-using viruses ^15^.

Receptor recognition of CoVs is usually species-specific and thus acts as a primary interspecies barrier at the entry level ^23^. Human emergence can occur upon host jumping and adaptive antigenic drift of coronaviruses ^24,25^. The order *Chiroptera* comprises more than 1,400 bat species with remarkable genetic diversity and wide geographic distribution. Bats are hosts of hundreds of known α- and β-CoVs and are important for viral evolution ^6^. ACE2 orthologs are largely conserved across mammalian species, while critical residues in viral receptor orthologs responsible for spike protein binding exhibit accelerated evolution in bats ^26,27^, resulting in species-specificity in supporting coronavirus binding and entry, as reported in SARS-CoV, SARS-CoV-2, and MERS-CoV ^28–30^. Of note, SARS-CoV-2 exhibits a broad host tropism with varying efficiency in using ACE2 orthologs from different bats and mammals ^26,31–33^. Adaptive mutations of the RBD region can occur when circulating in humans and other hosts ^34–36^.

We have previously shown NeoCoV and PDF-2180 selectively preferred ACE2 orthologs from Yangochiropteran bats (Yang-bats) for entry, whereas spike mutations like T510F on receptor binding motif (RBM) markedly enhanced hACE2 binding affinity ^15^. So far, the molecular basis of species-specific ACE2 recognition and potential host range of NeoCoV and PDF-2180 in diverse mammalian species remains unknown, which limited the assessment of the zoonotic risks of these viruses. By extensively examining the ACE2 orthologs from 102 mammalian species, we here demonstrated that NeoCoV and PDF-2180 can recognize ACE2 from a wide range of species and identified several critical host range determinants. We also showed that the cross-species transmission ability of these viruses could be further expanded through RBM mutations. Our data indicated a potentially broad host tropism of ACE2-using merbecoviruses, underscoring the necessity of viral surveillance in bats and other susceptible hosts to prevent future outbreaks.

## Results

### ACE2 orthologs from most Yin-bats are not well recognized by NeoCoV and PDF-2180

We previously tested a 293T stable cell library expressing 46 bat ACE2 orthologs and showed that ACE2 from most tested (14/15) Yin-bats were deficient in supporting NeoCoV and PDF-2180 RBD binding and pseudovirus entry ^15^. In this study, we re-examined the entry of NeoCoV and PDF-2180 mediated by 49 bat ACE2 orthologs stably expressed in 293T cells, including three new ACE2 orthologs from Yin-bats (Rcor, Raff, and Rmal) (**Figure 1A; Figure S1**). Consistent with our previous result, NeoCoV and PDF-2180 showed similar receptor usage profiles and were both less capable of using ACE2 from most (14/18) Yin-bats. However, the three newly-included Yin-bats’ ACE2 were functional, further suggesting that not all ACE2 from Yin-Bats were deficient in mediating viral entry (**Figure 1A**). Next, NeoCoV RBD binding and entry efficiency supported by Yin-Bats’ ACE2 were verified by 293T cells transiently expressing the 18 Yin-bats’ ACE2, with ACE2 orthologs from four Yang-bats (Nlep, Ppip, Lbor, and Nhum) and hACE2 as controls (**Figure 1B and 1C**). Immunofluorescence assay detecting the C-terminal fused 3×Flag tags indicated these receptors were expressed at a similar level (**Figure S2**). We next tested whether the suborder-specific bat ACE2 preference can also be observed in other representative ACE2 using viruses with distinct receptor binding modes (**Figure 1D**). In line with the different RBD binding footprints on ACE2, SARS-CoV-2 and NL63 exhibited very distinct receptor usage profiles among the 22 tested bat ACE2 orthologs as indicated by the RBD binding and pseudovirus entry assays **(Figure 1E; Figure S2)**. SARS-CoV-2, which is supposed to share common ancestors infecting Yin-bats (*Rhinolophus malayanus*, Rmal) ^8^, can efficiently use 13 (>10% to hACE2, Hsap) ACE2 orthologs from Yin-bats. Although NL63 relatives were sampled in Yang-bat (*Perimyotis subflavus*)^37^, most (14/18) (>10% to hACE2, Hsap) tested Yin-bats’ ACE2 can support efficient binding and entry of NL63. These data indicate that the suborder-specific ACE2 usage is not strictly consistent with the phylogeny of their natural hosts, and there are specific host range determinants to be identified.

**Figure 1.**
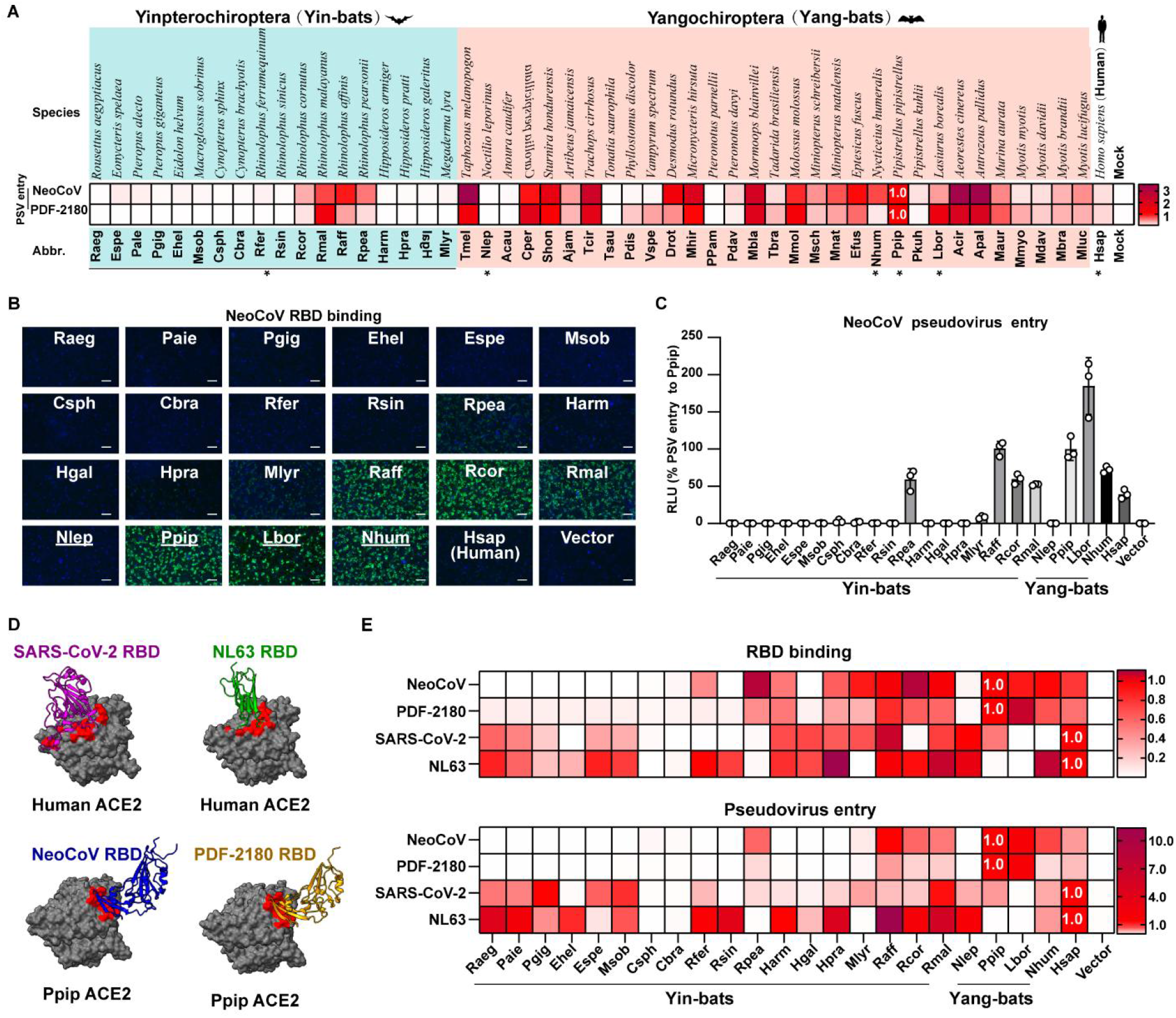
Species-specific ACE2 usage of NeoCoV, PDF-2180, and other ACE2-using viruses. (**A**) Heat map of pseudovirus entry efficiency (RLU relative to RLUPpip) of NeoCoV and PDF-2180 on HEK293T cells stably expressing 49 bat ACE2 orthologs. * indicated ACE2 selected for subsequent characterizations. Upper, species names. Lower, abbreviation of species names. Yinpterochiroptera (Yin-bats) and Yangchiroptera (Yang-bats) are indicated with cyan and pink backgrounds. The pseudovirus entry efficiency mediated by Ppip ACE2 was set as 1.0. (**B-C**) Most ACE2 orthologs from Yin-bats were deficient in supporting NeoCoV and PDF-2180 binding and pseudovirus entry. NeoCoV RBD-hFc binding (**B**) and pseudoviruses entry efficiency (**C**) were evaluated on HEK293T cells transiently expressing the indicated ACE2 orthologs. Vector plasmid was used as a negative control. The underlines in **B** indicate species from Yang-bats. Scale bar in **B**:100 μm. (**D**) Distinct RBD binding modes of four ACE2 using coronaviruses. RBD footprints of SARS-CoV-2 (PDB: 6M0J, purple), NL63 (PDB: 3KBH, green), NeoCoV (PDB: 7WPO, blue) and PDF-2180 (PDB: 7WPZ, yellow) on indicated ACE2 were highlighted in red. **(E)** Heat map of the RBD binding (RFU) and pseudoviruses entry efficiency (RLU) of ACE2 using coronaviruses on HEK293T cells transiently expressing the indicated ACE2 orthologs. The binding and pseudoviruses entry efficiency mediated by Ppip ACE2 was set as 1.0 for NeoCoV/PDF2180, the values of hACE2 were set as 1.0 for SARS-CoV-2 and NL63. Data are presented as mean ± SD for n=3 biologically independent cells for **C**. Data are presented as mean for n=3 biologically independent cells for **A** and **E**. Data representative of two independent experiments. RFU: relative fluorescence unit. RLU: relative light units.

### NeoCoV and PDF-2180 exhibit a broad receptor tropism across non-bat mammals

We further explored the ability of ACE2 orthologs from 53 non-bat mammals, most of which were selected and tested for SARS-CoV-2 in a previous study ^26^. These species include wild and domestic animals, and common animals in zoos and aquaria that belong to ten mammalian orders: Carnivora, Primates, Artiodactyla, Rodentia, Cetacea, Perissodactyla, Diprotodontia, Pholidota, Erinaceomorpha, and Lagomorpha (**Figure S1**). These animals are either in frequent contact with humans, used as model animals, endangered animals, or potential natural or intermediate hosts of CoVs. The expression of all ACE2 orthologs was verified by immunofluorescence (**Figure S3**). We then conducted NeoCoV and PDF-2180 RBD binding and pseudovirus entry assays to test their receptor function by transiently expressing them in 293T cells, with ACE2 from Ppip bat (Ppip ACE2) as a positive control. The binding and entry assays showed generally consistent results in most species (**Figure 2A, S3**). 47 out of 53 ACE2 orthologs could support entry of both viruses, albeit with various efficiencies (20 to 100% to Ppip ACE2 for NeoCoV). Overall, the several primates showed relatively low efficiency to support RBD binding and entry of both viruses, including humans. The six ACE2 orthologs showing undetected or very limited entry (<20% to Ppip ACE2 for NeoCoV) were from five different orders: Sape (*Sapajus apella*), Sbol (*Saimiri boliviensis*), Sscr (*Sus scrofa*), Nasi (*Neophocaena asiaeorientalis*), Csim (*Ceratotherium simum*), and Pcin (*Phascolarctos cinereus*) (**Figure 2A**). Representative RBD binding and entry efficiency of NeoCoV were shown (**Figure 2B and 2C**). Next, we demonstrated the NeoCoV RBD binding and entry efficiency with ACE2 orthologs from the above species and with seven CoV host-related species as controls^20,25,38–40^. The expression levels of the selected ACE2 orthologs were verified by Western blot (**Figure 2D and E**). As expected, the results confirmed that the ACE2 from seven CoV host-related species supported efficient pseudovirus entry compared to the six deficient ACE2 orthologs. (**Figure 2F and G**). Collectively, these data demonstrate that ACE2 orthologs from a wide range of species can act as functional receptors for NeoCoV and PDF-2180, suggesting these species might be susceptible to the infection of these viruses.

**Figure 2.**
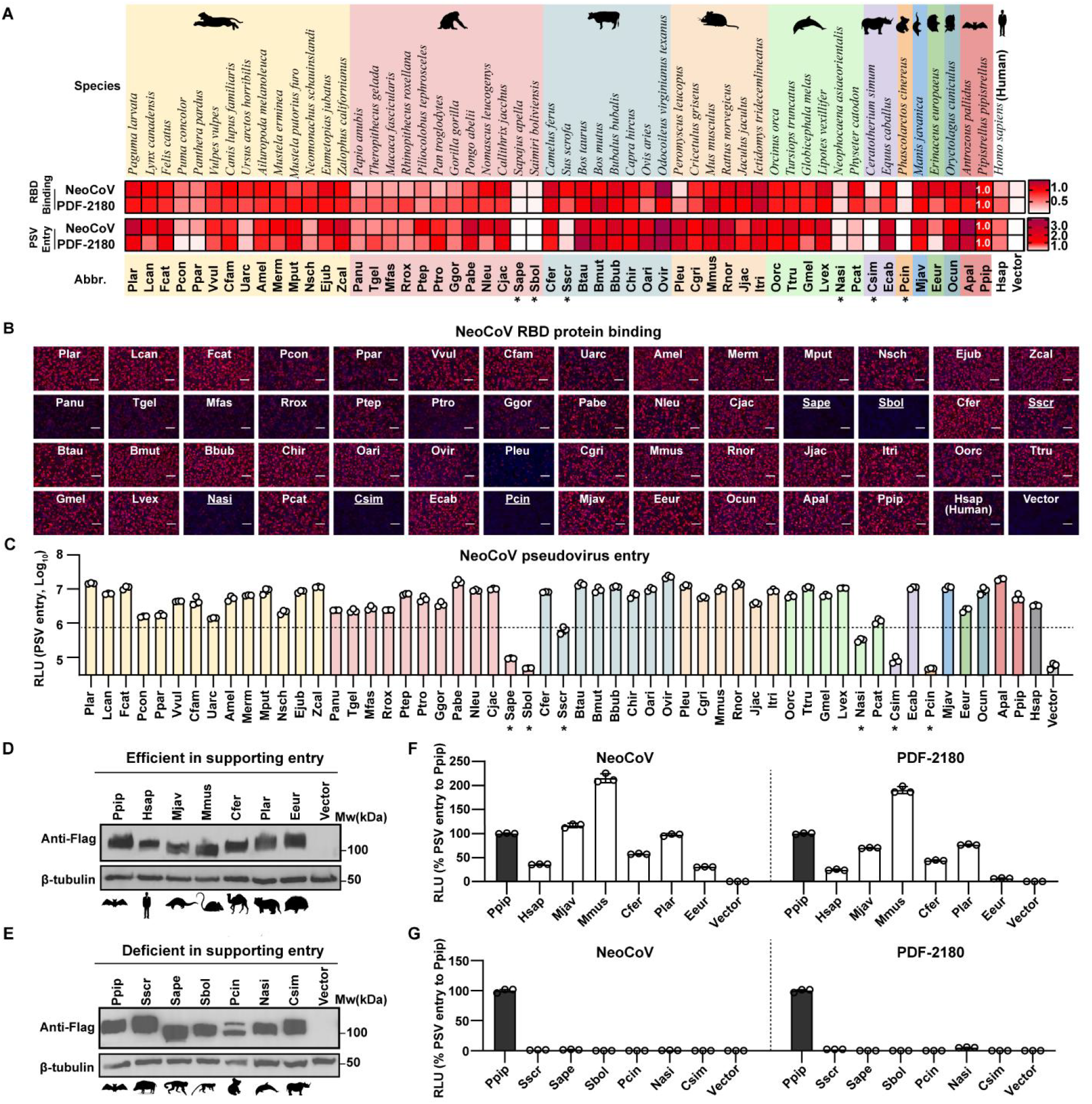
NeoCoV and PDF-2180 recognize a wide range of mammalian ACE2 orthologs. (**A**) The heat map of RBD binding (RFU relative to RFUPpip) and pseudoviruses entry efficiency (RLU relative to RLU_Ppip_) of NeoCoV and PDF-2180 mediated by various non-bat mammalian ACE2 orthologs. The pseudovirus entry efficiency and RBD binding on Ppip ACE2 were set as 1.0. *: RLU<20% RLU_Ppip_. Upper: species name. Lower: abbreviation of species name. Mammals belonging to different orders are indicated with colored backgrounds, from left to right: Carnivora, Primates, Artiodactyla, Rodentia, Cetacea, Perissodactyla, Diprotodontia, Pholidota, Erinaceomorpha, and Lagomorpha. **(B)** Binding of NeoCoV RBD-hFc with HEK293T transiently expressing mammalian ACE2 cells analyzed by immunofluorescence assay detecting the hFc. Scale bar: 100 μm. Underline indicate the six non-supportive ACE2 orthologs. **(C)** Entry efficiency of NeoCoV pseudoviruses in HEK293T cells transiently expressing the indicated mammalian ACE2. Dashed line: 20% RLU_Ppip_. *: RLU<20% RLU_Ppip_. **(D-E)** Western blot showing the expression of ACE2 orthologs from selected CoV host-related species (**D**) or NeoCoV/PDF-2180 non-supportive species (**E**) in HEK293T cells. **(F-G)** Pseudoviruses entry efficiency of NeoCoV and PDF-2180 in HEK293T cells expressing the indicated ACE2 orthologs. Data are presented as mean ± SD for n=3 biologically independent cells for C. Data are presented as mean ± SEM for n=3 biologically independent cells for F and G. Data representative of two independent experiments for A-G. RLU: relative light unit. Mw: molecular weight.

### Identification of host tropism determinants restricting NeoCoV and PDF-2180 recognition

We next sought to identify the host tropism determinants through comparative analyses focusing on the two critical RBD interacting loops on ACE2 orthologs, each containing a glycosylation site in Ppip ACE2 **(Figure 3A)**. We start with the bat species, considering the numbers and sequence variations in the non-supportive ACE2 orthologs from Yin-bats. We first conducted multi-sequence alignment and conservation analyses based on 49 bat ACE2 sequences (from 18 Yin- and 31 Yang-bats), which are separated into two groups by their ability to support NeoCoV entry **(Figure 3B; Figure S1 and S4)**. The bat ACE2 orthologs are largely conserved, while highly variable residues can be found within the viral binding loops. The comparative analysis highlighted four candidate determinants (from A-D) with contrasting residue frequencies across the two groups, all located in the receptor binding interface **(Figure 3C)**. According to the presence of putatively deficient/sub-optional residues, we defined and classified the ACE2 orthologs deficient in viral receptor function into several defect types **(Figure 3D).** For example, Raeg ACE2 was considered defect type ABCD as it carries putatively deficient residues on all four determinants. We next analyzed the impact of the deficient/sub-optimal ACE2 residues on viral RBD recognition based on the structure of the Ppip ACE2-NeoCoV-RBD complex **(Figure 3E-I)**. Glycosylations sites (N-X-T/S) in A and C that are required for the glycan-protein interactions are absent in several Yin-bats’ ACE2. Determinant A glycosylation (N54 glycosylation) (**Figure 3F**) is more important than determinant C glycosylation (N329-glycosylation) (**Figure 3H**) for viral receptor interaction as it mediated a critical protein-glycan interaction underpinning the RBD binding away from the main protein-protein binding interface. Comparatively, determinant C glycosylation only partially contributes to the extensive interactions around this region. Residues in determinants B (**Figure 3G**) and D (**Figure 3I**) of Yin-bats’ ACE2 either abolish the polar contacts (e.g., two salt bridges formed by residues E305 and D338, respectively) or introduce steric hindrance (e.g., E305K) that reduce the binding affinity. It is worth mentioning that determinant D, especially the D338, is also a critical host range determinant restricting hACE2 from efficiently supporting the binding and entry of the two viruses ^15^. A similar comparative analysis was also conducted based on ACE2 orthologs from non-bat mammals, which is less informative than bat species as only six non-bat mammalian species are potentially non-permissive **(Figure S5)**.We conducted sequence conservation analysis of non-bat ACE2 orthologs based on three groups: the total 53 non-bat mammals, six deficient ACE2 orthologs, and 30 competent ACE2 orthologs (NeoCoV entry efficiency > Ppip ACE2), respectively (**Figure 2C**). Compared with the four determinants identified among bat species, the analysis of ACE2 orthologs from the six non-bat mammals mainly pointed to putatively deficient /sub-optional residues in determinant B (residue 305), which is associated with a loss of a salt bridge interaction as indicated by the cryo-EM structure of NeoCoV RBD-Ppip ACE2 complex (**Figure 3G**) ^15^.

**Figure 3.**
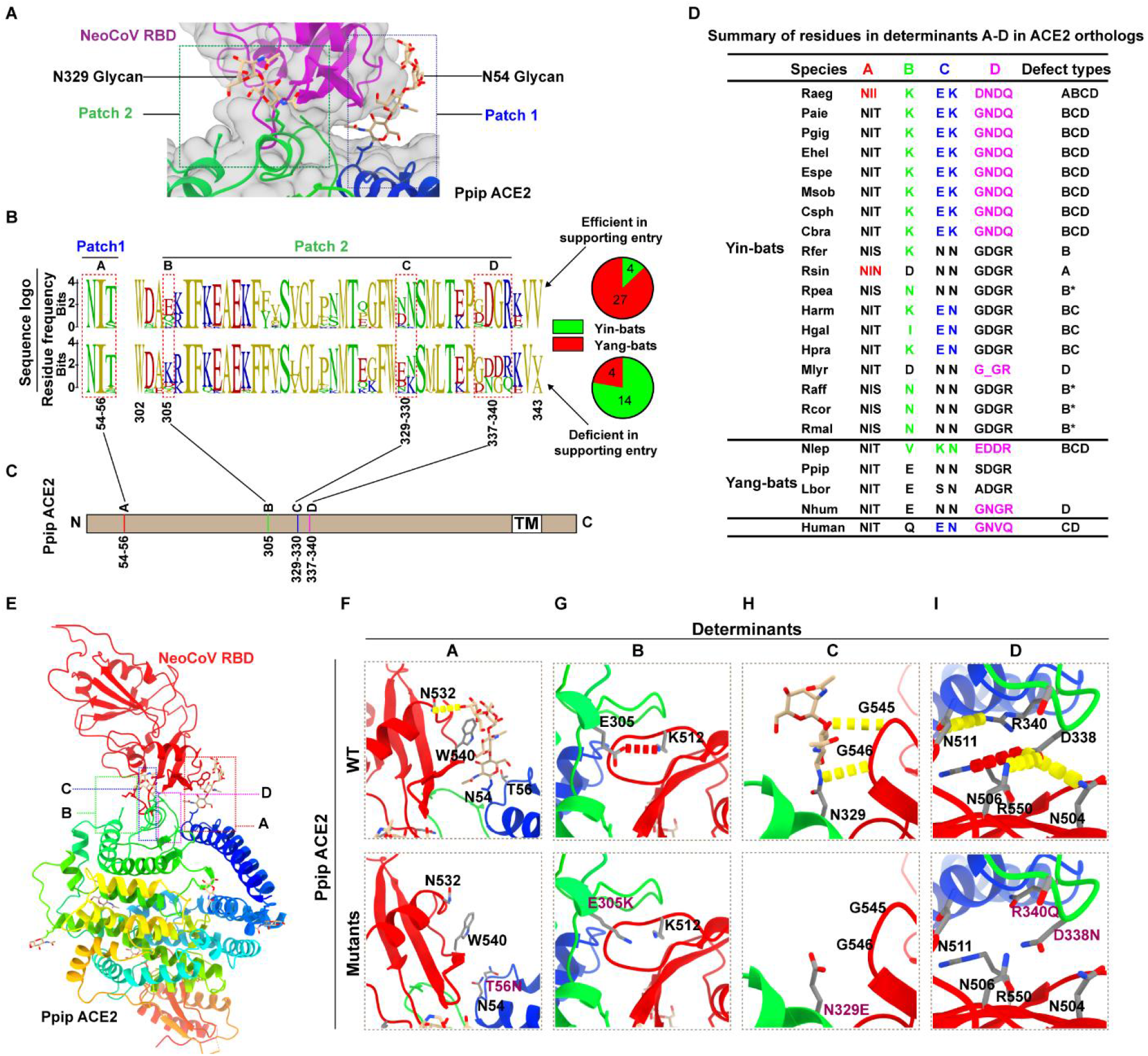
Identification of host range determinants restricting NeoCoV and PDF-2180 recognition. (**A**) Magnified view of the binding interface of NeoCoV RBD (purple) and Ppip ACE2 (rainbow). Patch 1 and patch 2 indicate two main interaction regions, each containing a glycosylation on Ppip ACE2. (**B-C**) Comparative sequence analysis predicting the potential host range determinants. (**B**) Residue conservation of the two critical viral binding loops based on sequences of 49 ACE2 orthologs, which are separated into two groups according to their entry-supporting efficiency. Upper: efficient in supporting NeoCoV and PDF-2180 entry (>10% RLU_Ppip_). Lower: deficient in supporting NeoCoV and PDF-2180 entry (<10% RLU_Ppip_). The pie charts summarized the numbers of Yin-bats and Yang-bats in each group. (**C**) The four variable regions showing group-specific residue frequencies were defined as determinants A-D. Sequence numbers were based on Ppip ACE2. TM, transmembrane motif. (**D**) Summary of defect types of bat and human ACE2 orthologs according to their residue features in determinants A-D. The predicted sub-optional residues in determinants A (54-56), B (305), C (329-330), and D (337-340) were highlighted with red, green, blue, and magenta, respectively. B*, sub-optional but acceptable residues for hydrogen bond formation. (**E-I**) Structural analyses of the impact of sub-optional residue substitution on the interaction between Ppip ACE2 and NeoCoV RBD. (**E**) Structure of Ppip ACE2 and NeoCoV RBD complex, with each determinant indicated by dashed boxes. (**F-I**) Magnified view of the interface of determinants A-D according to the WT (upper) and mutated (lower) Ppip ACE2, respectively. All structures are shown as ribbon representations, with key residues rendered as sticks. Salt bridges and hydrogen bonds are shown as red and yellow dashed lines. Mutated residues were highlighted in purple.

### Functional verification of host tropism determinants of NeoCoV and PDF-2180 in bats

To verify the predicted determinants, we picked representative ACE2 orthologs of specific defect types for mutagenesis assays to improve their receptor recognition. Specifically, We generated a series of gain of function mutations based on ACE2 orthologs from Rsin (type A), Rfer and Rpea (type B), Hgal (type BC), Nlep (type BCD), and Raeg (type ABCD). The results showed that the N55T point mutation, which introduces an N53 glycosylation site, markedly improved the receptor function of Rsin ACE2 (type A) **(Figure 4A and 4B; Figure S6)**. Rfer-K305E (type B) and Rpea-N305E (type B*) also showed significantly improved receptor function after introducing the optimal residue for potential salt bridge formation (**Figure 4C and 4D; Figure S6**). Efficient binding and pseudovirus entry mediated by bat ACE2 mutants from Hgal (type BC) and Nlep (type BCD) are achieved after corresponding residues are replaced by Ppip ACE2 counterparts **(Figure S6)**. Remarkably, a gradual gain of receptor function of Raeg ACE2 (type ABCD) can be observed following stepwise increased substitutions of determinants A, AB, ABC, and ABCD with Ppip ACE2 counterparts **(Figure 4F and 4G)**. We further evaluated the binding affinities between viral RBD proteins and representative WT and mutant ACE2 by Flow cytometry and Bio-layer interferometry (BLI) assays **(Figure 4H and 4I)**. As expected, the binding affinities between WT Raeg/Rsin ACE2 and NeoCoV/PDF-2180 RBDs were below the detection limit **(Figure S7)**, while corresponding mutations significantly improved RBD binding efficiency, with apparent binding affinities (K_D, app_) ranging from 6.94×10^-10^ to 2.35×10^-8^ **(Figure 4H and 4I)**. Restored receptor function of Rsin ACE2-N55T and Raeg ACE2-ABCD was also confirmed in a bat cell line Tb 1 Lu **(Figure 4J; Figure S7)**. Together, our results demonstrated that deficient/sub-optional residues in the four predicted host range determinants restricted the recognition of most bat ACE2 orthologs by NeoCoV and PDF-2180.

**Figure 4.**
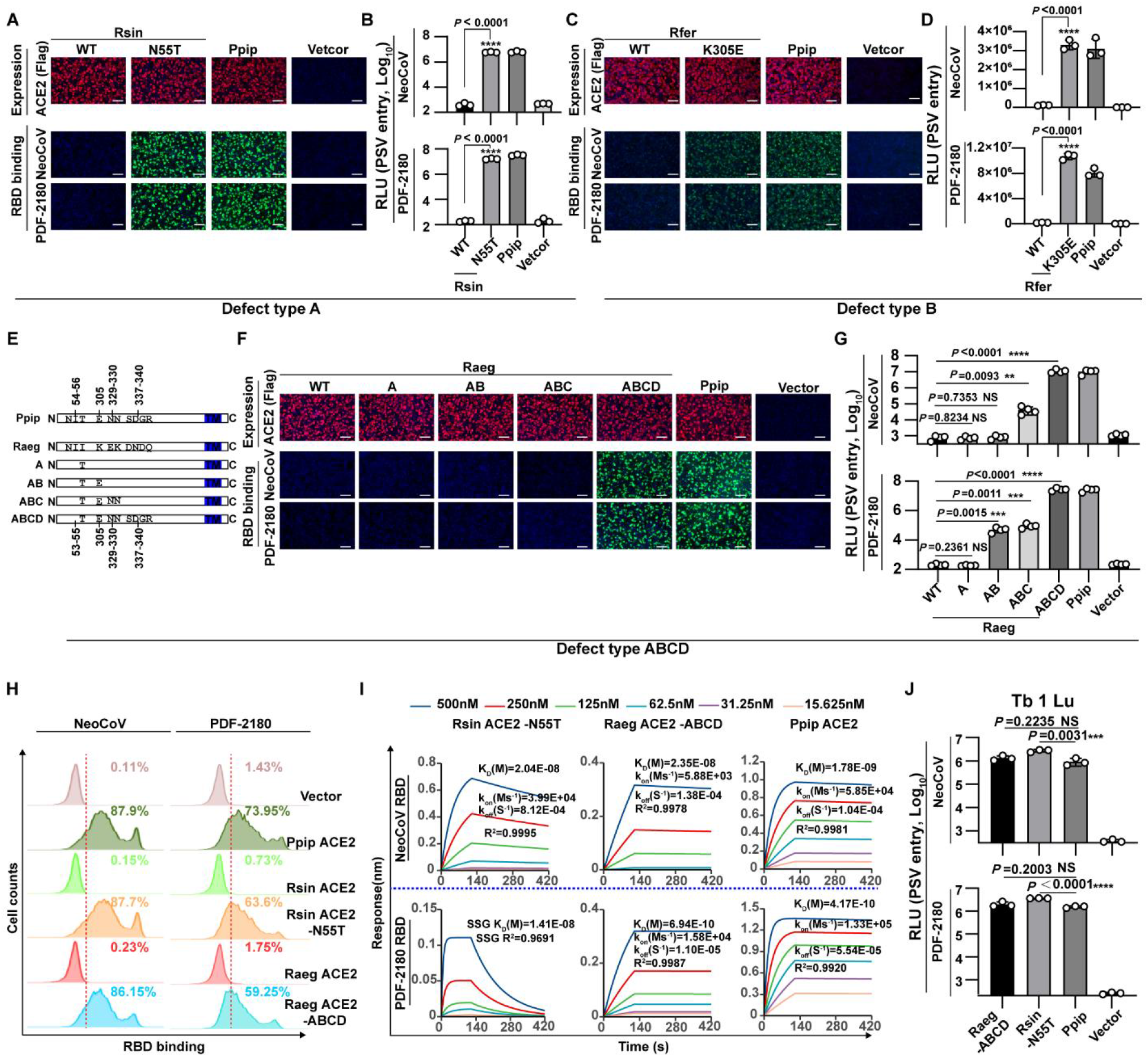
Verification of determinants crucial for species-specific receptor usage of NeoCoV and PDF-2180 in bats. (**A-G**) Gain of receptor function of Yin-bats’ ACE2 orthologs in supporting NeoCoV and PDF-2180 RBD binding (**A, C, F**) and pseudoviruses entry (**B, D, G**) through the indicated mutations. **(A-B)** Rsin ACE2 (defect type A); (**C-D**) Rfer ACE2 (defect type B); (**E-G**) Raeg ACE2 (defect type ABCD). (**E**) Schematic illustration of Raeg ACE2 swap mutants carrying the indicated Ppip ACE2 counterparts. Data related to defect type B*, BC, and BCD can be found in Figure S6. N: N-terminus, C: C-terminus. (**H**) Flow cytometry analysis of NeoCoV and PDF-2180 RBD-hFc binding with HEK293T cells transiently expressing the indicated ACE2 orthologs. The vector was used as a negative control. The red dashed lines indicate the threshold to define positive cells. (**I**) BLI assays analyzing the binding kinetics between NeoCoV RBD-hFc/PDF-2180 RBD-hFc and the indicated ACE2 ectodomain proteins with indicated mutations. (**J**) NeoCoV and PDF-2180 pseudoviruses entry in Tb1 Lu cells transiently expressing the indicated ACE2 orthologs at 16 h post-infection. Data are presented as mean ± SD for n=3 biologically independent cells for **B** and **D**, n=4 for **G**. Data representative of three independent experiments for A-G. Representative data of three independent experiments are presented as mean for n=3 technical repeats for H. Representative data of two independent experiments are presented as mean ± SD for n=3 biologically independent cells for J. Two-tailed unpaired Student’s *t*-test for **B**, **D**, **G**, and **J**; * p<0.05, ** p <0.01, *** p <0.005, and **** p <0.001. NS: not significant. RLU: Relative luciferase unit. Scale bar in A, C, and F: 100 μm.

### Genetic determinants restricting NeoCoV and PDF-2180 entry in non-bat mammals

We next explored the genetic determinant restricting NeoCoV/PDF-2180 recognition from ACE2 orthologs from the six non-bat mammals: Pig (Sscr), Koala (Pcin), two closely related New World monkeys (Sape, Sbol), and two endangered animals Finless Porpoise (Nasi) and Southern white rhinoceroses (Csim). In line with the previous hypothesis that site 305 is a crucial host range determinant, none of the six deficient ACE2 orthologs carries an E at this site for optimal salt bridge formation. Therefore, we generated 305E mutants (replacing the residue of 305 to E) for the six ACE2 orthologs and tested their ACE2 function. All these mutants were well expressed and the 305E mutation rendered efficient RBD binding for ACE2 orthologs from Sscr, Nasi, and Csim, but not for ACE2 orthologs from Sape, Sbol, and Pcin (**Figure 5A**). NeoCoV and PDF-2180 pseudovirus entry assay showed largely consistent results with the RBD binding assay (**Figure 5B**).

**Figure 5.**
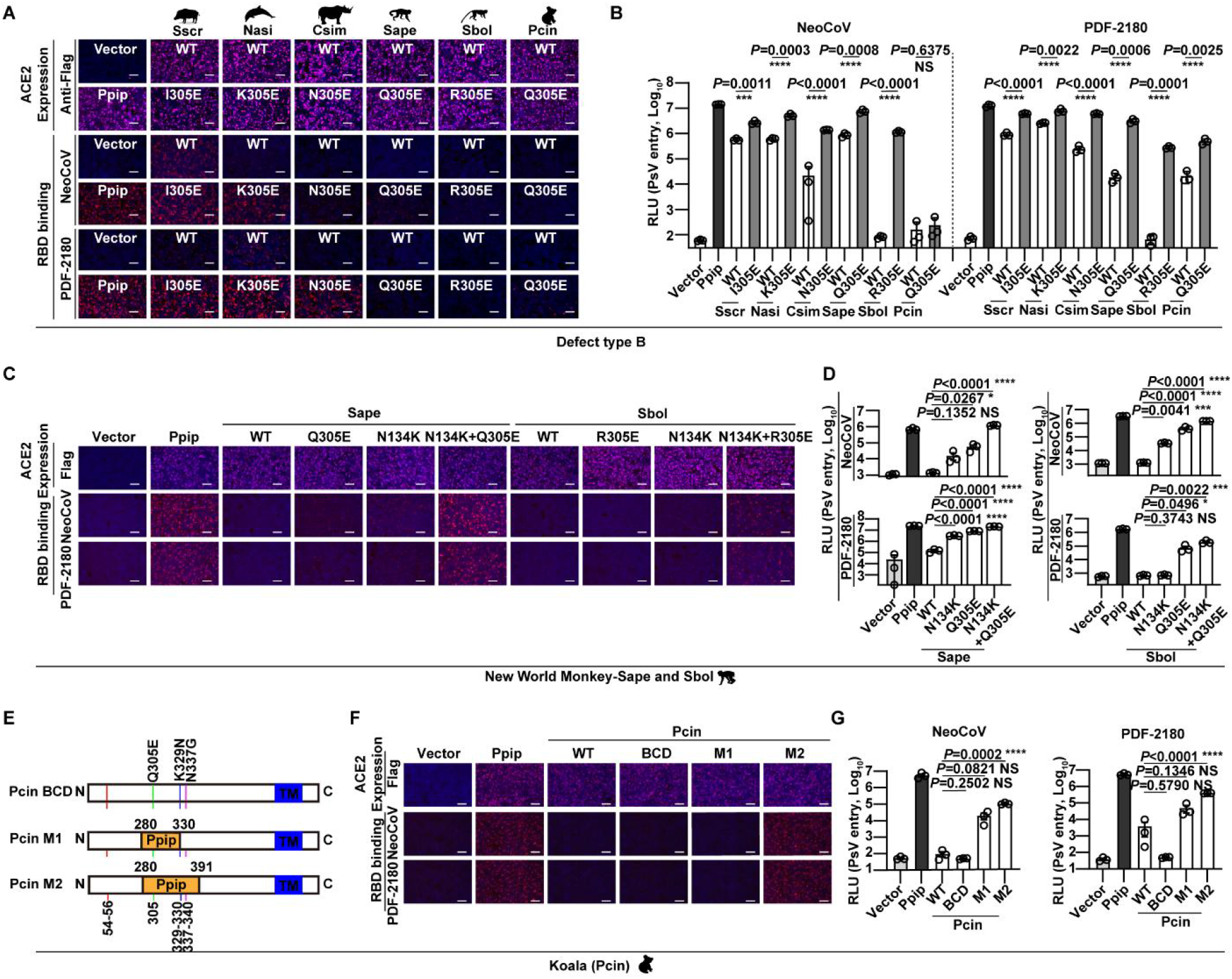
Determinants restricting the recognition of NeoCoV and PDF-2180 of ACE2 orthologs from non-bat mammals. (**A-G**) Gain of receptor function of indicated mammalian ACE2 orthologs in supporting NeoCoV and PDF-2180 RBD binding (**A**, **C**, **F**) and pseudoviruses entry (**B**, **D**, **G**) through the indicated mutations. (**A-B**) The six non-supportive mammalian ACE2 (defect type B); (**C-D**) Sape and Sbol ACE2 (defect type B); (**E-G**) Pcin ACE2 (defect type BCD). (**E**) Schematic illustration of Pcin ACE2 mutants carrying substitutions of Ppip counterparts. Data are presented as mean ± SD for n=3 biologically independent cells for **B** and **D.** Data are presented as mean ± SEM for n=3 biologically independent cells for **B, D,**and **G.**Data representative of two independent experiments for **A-G**. Two-tailed unpaired Student’s t-test; * p<0.05, ** p <0.01, ***p <0.005, and **** p <0.001. RLU: relative light unit. Scale bar represents 100 μm for **A**, **C,** and **F**.

Since the 305E mutation alone is insufficient to fully recover receptor function for Sape, Sbol, and Pcin ACE2, we further explore other genetic determinants restricting their recognition. For Sape and Sbol ACE2, we made chimeric ACE2 with specific regions substituted by the phylogenetically close-related Cjac ACE2. The result showed that a significant gain of receptor function of Sape could be observed when aa1-251 or aa125-251 were replaced by Cjac ACE2 counterparts. Fine mapping of region aa125-251 targeted residue 134 as a specific genetic determinant for Sape and Sbol (**Figure S8**). A better gain of receptor function of Sape and Sbol ACE2 orthologs can be observed upon E134K and Q/R305E double mutation. (**Figure 5C and 5D**). Koala (Pcin) ACE2 is phylogenetically distant to ACE2 from other mammals (**Figure S1**). A previous study reported that T31K and F83Y double mutations work in cooperation to restore receptor function of Koala ACE2 to support SARS-CoV-2 entry ^32^. However, the Pcin ACE2 with BCD substitutions remains defective in supporting NeoCoV and PDF-2180 binding and entry (**Figure 5E-G)**. We then generated ACE2 chimera with specific regions replaced by PpipACE2 equivalent sequences (**Figure 5E)**. Our result showed that a significant gain of receptor function could be observed on mutant 2 (aa280-391) but not on mutant 1 (aa280-330), suggesting a host range determinant between aa330-391 (**Figure 5F and 5G)**. Together, our results highlighted the critical role of site 305 for host tropism determination in mammals albeit the presents of other species-specific determinants.

### RBM mutations further expand the potential host range of NeoCoV

We previously showed that specific mutations in NeoCoV and PDF-2180 RBM confer more efficient hACE2 recognition ^15^. For NeoCoV, the substitution of its T510 by the PDF-2180 equivalent residue F (T510F) with higher hydrophobicity enhanced its interaction with a hydrophobic pocket of human ACE2 (**Figure 6A**). Sequence analysis of the 102 ACE2 orthologs tested in this study indicates residues constituting this hydrophobic pocket are highly conserved among mammals (**Figure 6B**). Thus, we hypothesized that mutations such as T510F could expand the potential host range by enhancing hydrophobic interactions with this highly conserved pocket. As expected, NeoCoV-T510F efficiently binds with most tested ACE2 orthologs and can achieve efficient entry by these receptors, except for Rsin and Raeg ACE2 lacking the critical N53 glycosylations (**Figure 6C-D and Figure S4**). Further experiments demonstrated that Rsin ACE2-N55T and Raeg ACE2-I55T with functional N53 glycosylation sites also achieved efficient NeoCoV-T510F entry (**Figure 4E-F**). In addition, the NeoCoV-T510F also showed an improved ability to recognize ACE2 from non-bat mammals, as indicated by their increased RBD binding and pseudovirus entry efficiency with the six ACE2 orthologs that are not recognized by the WT viruses (**Figure 6G and 6H**). These results indicate that NeoCoV, PDF-2180, or related viruses may expand their potential host range to Yin-bats and other non-permissive mammals, including humans, through antigenic drifts on RBM, such as T510F.

**Figure 6.**
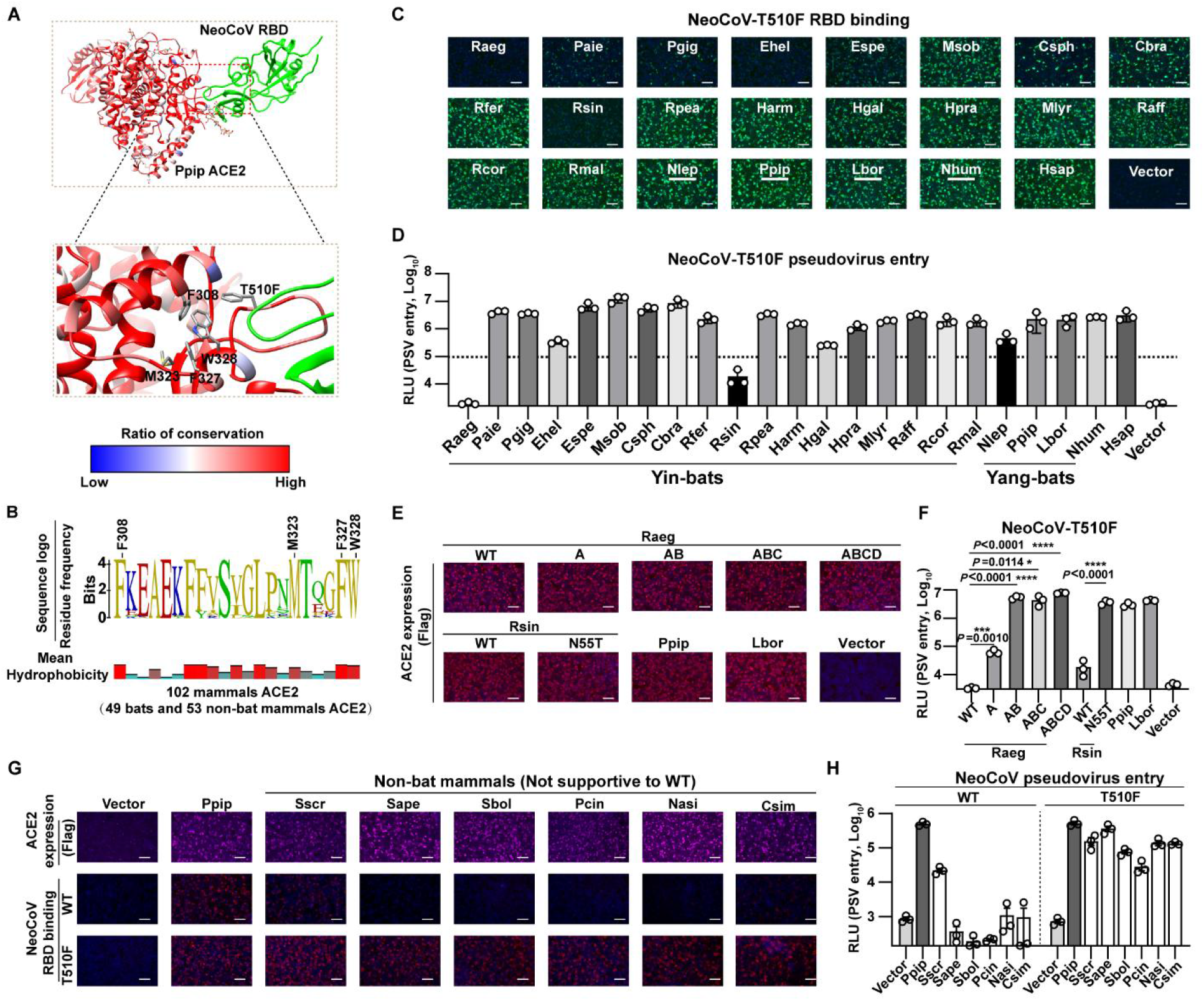
RBM mutation T510F further expanded the potential host range of NeoCoV. (**A**) Structure showing the interaction interface between NeoCoV-T510F and a conserved hydrophobic pocket in Ppip ACE2. The side chains of hydrophobic residues constituting the pocket were indicated as sticks in the magnified view. The blue and red colors of the Ppip ACE2 represent the conservation ratio based on ACE2 orthologs from 49 bats and 53 non-bat mammals. (**B**) The sequence conservation of residues surrounding the conserved hydrophobic pocket based on ACE2 orthologs from 49 bats and 53 non-bat mammals. The residues that constitute the pocket were indicated with sequence numbers. The side chain hydrophobicity is shown below. (**C-D**) NeoCoV RBD-T510F-hFc RBD binding (**C**) and pseudovirus entry (**D**) in HEK293T cells transiently expressing the indicated bat ACE2 orthologs. (**E**) The immunofluorescence analyzing the expression level of Raeg and Rsin ACE2 orthologs and their mutants. (**F**) The NeoCoV-T510F pseudovirus entry efficiency mediated by the indicated ACE2. (**G-H**) Efficiency of NeoCoV-T510F RBD binding (**G**) and pseudovirus entry (**H**) on HEK293T cells transiently expressing the indicated mammalian ACE2 orthologs. Data are presented as mean ± SD for n=3 biologically independent cells for **D** and **F**. Data are presented as mean ± SEM for n=3 biologically independent cells for **H.** Data representative of two independent experiments for **C-H**. Two-tailed unpaired Student’s *t*-test; * p<0.05, ** p <0.01, *** p <0.005, and **** p <0.001. RLU: relative light unit. Scale bar represents 100 μm for **C, E,** and **G**.

## Discussion

Global transmission of coronaviruses with higher pathogenicity, like MERS-CoV, could be more devastating than the COVID-19 pandemic ^41^. Up to August 2022, MERS-CoV caused 2591 Laboratory-confirmed cases and 894 death worldwide since its emergence in Saudi Arabia in April 2012 ^1–11^

Fortunately, MERS-CoV appears to have relatively low transmissibility with a reproductive number (R0) around 0.69, which result in a gradual reduction of infected cases since 2016^42 11^. Whether the relatively low transmission rate is associated with the DPP4 receptor usage or the unaccomplished human adaptation remains an open question. Yet, the zoonotic emergence of MERS-CoV-related coronaviruses may occur and even develope into a pandemic. The origin of MERS-CoV remains a mystery, while hypotheses have been proposed that MERS-CoV may arise from the recombination and evolution of MERS-related bat coronaviruses, such as HKU4, NeoCoV, and PDF-2180^12,43–46^.

Phylogenetically distant coronaviruses evolved to use ACE2 as their common receptors ^19^. To date, coronaviruses of three different subgenera have evolved to engage ACE2 for cellular entry, including NL63 (Setracovirus subgenus, α-CoV), many SARS-related CoVs (*Sarbecovirus* subgenus, β-CoV), and the two recently reported MERS-related CoVs (*Merbecovirus* subgenus, β-CoV) in this study. The distinct viral RBD structure and ACE2 binding footprints suggest convergent evolutionary histories of receptor acquisition and adaptation of these viruses. The reason for ACE2 preference among coronaviruses remains unknown. However, it should be noted that ACE2 holds the potential to be utilized by coronaviruses to achieve efficient airborne transmission, considering the highly transmissible SARS-CoV-2 Omicron variant ^47^ So far, structures very similar to NeoCoV and PDF-2180 RBDs were not reported in other bat coronaviruses, and the closest RBD homologs were found in hedgehog merbecoviruses which do not recognize ACE2 ^15,20,22^. Thus, knowledge of the transmission ability and host tropism of NeoCoV and PDF-2180, the only two known ACE2-using coronaviruses to date, is crucial for assessing the zoonotic risk of these viruses.

We demonstrated that NeoCoV and PDF-2180 could efficiently use most ACE2 orthologs from 102 mammalian species across 11 orders, highlighting a potentially board host tropism of ACE2-using merbecoviruses. It showed that these viruses displayed a bat-specific phenotype preferring ACE2 orthologs from Yang-bats but not from most Yin-bats, which is not observed in NL63 and SARS-CoV-2. This bat ACE2 preference is in line with the observation that most merbecoviruses were sampled in bats belonging to the family *Vespertilionidae* (vesper bats), the largest family of Yangochiroptera (**Figure S1**), including the hosts of NeoCoV and PDF-2180 ^6^. Comparatively, most sarbecoviruses were identified in Yin-bats, particularly *Rhinolophine* or *Hzpposideros*^6,48^. The differences in host preference likely limited the opportunities for cross-lineage recombination of these high-risk viruses. Only six tested non-bat mammals from 5 different orders were found to be almost non-supportive, while human ACE2 exhibited a relatively weak receptor function among the tested ACE2 orthologs.

We here revealed that specific residues in the viral binding interface determine ACE2 tropism of merbecoviruses. Interestingly, glycan-protein interactions play a crucial role in ACE2 recognition of merbecoviruses, especially the crucial interaction mediated by N54-(or N53 in Rsin and Raeg) glycosylation (determinant A). Similar glycan-protein interaction in receptor engagement has also been reported in other coronaviruses, including SARS-CoV-2 and human-infecting CCoV-HuPn-2018 ^6,48–50^. It could be interesting to investigate their contribution to host tropism in other coronaviruses. Another glycan-related determinant is related to the N329 (or N330 in some species) glycosylation. As only some ACE2 orthologs from Yang-bats are glycosylated at this site, its contribution to binding affinity and species specificity is less prominent than the N54 glycosylation, probably due to the compensation of nearby protein-protein interactions, while it has to be noted that all tested ACE2 orthologs from non-bat mammals carry the N54-glycosylation. Besides the two glycosylation related determinants, determinants B and D participate in protein-protein interactions required for effective receptor recognition, especially salt bridges. Although determinant D has been demonstrated to restrict hACE2 recognition, the interaction mediated by determinant B (E305) plays a more important role in host range determinants in both bats and non-bat mammals via salt bridge formation. N305 in some ACE2 orthologs may form sub-optional hydrogen bonds with the viruses, while K305 seems unable to interact with the viruses and may even result in steric hindrance. In addition, additional determinants beyond the viral binding surface, like N134K in two New World primates, Sape and Sbol, also contribute to host tropism. The mechanism may involve their influence on the ACE2 structure that indirectly affects the viral binding, which could be elucidated by structural analysis in future studies. The molecular basis of why koala (Pcin) ACE2 could not be functional by modifying the characterized determinants remains unclear, while a full gain of function can be achieved through large fragment substitution, suggesting other critical genetic determinants are restricting Koala from NeoCoV infection that can be characterized in future studies.

Although the incompatible receptor recognition sets a primary barrier for inter-species transmission of coronaviruses, viruses could achieve host jumping via adaptive antigenic drift^51^. Here we show that the T510F mutation in the NeoCoV spike, increasing binding affinity via interacting with a conserved hydrophobic pocket, broadens the potential host range. Our results indicate that NeoCoV and related viruses hold the potential to break the current host range barrier via adaptive antigenic drift or recombination in bats or other mammals. It should be noted that host immune responses and other host factors required for viral infections also play important roles in receptor-independent host tropism determination ^52^. Thus, it remains unknown whether NeoCoV carrying T510F mutant, which has not been found in nature, can readily infect humans. However, there might be more ACE2-using merbecoviruses with better human ACE2 recognition that need to be added to our radar. Thus more attention should be paid to the surveillance of these viruses in the wild.

Together, we revealed a broad receptor usage of ACE2-using merbecoviruses across mammals and characterized the critical genetic determinants restricting the host range. Our study adds knowledge to the molecular basis of species-specific ACE2 recognition of merbecoviruses, highlighting the importance of in-depth research of these potentially high-risk viruses to prepare for potential future outbreaks.

## Acknowledgments

Huan Yan was supported by China NSFC projects (32270164, 32070160) and the Special Fund for COVID-19 Research of Wuhan University. We thank Beijing Taikang Yicai Foundation for their support of this work. We thank Qihui Wang (CAS Key Laboratory of Pathogenic Microbiology & Immunology, China) and Qiang Ding (Tsinghua University) for their kind offer of some mammalian ACE2 plasmids.

## Author contributions

Conceptualization, H.Y., C.B.M., and L.C.; methodology, C.B.M., L.C.,Q.X., M.X.G., L.L.S., C.L.W., J.Y.S., F.T., P.L, and M.L.H.; formal analysis, C.B.M., L.C., Q.X., and H.Y.; investigation, C.B.M., C.L., Q.X., and H.Y.; writing—original draft, H.Y., C.B.M., and L.C.; writing—review& editing, all authors; visualization, C.B.M. and L.C.; supervision and funding acquisition, H.Y.

## STAR METHODS

### KEY RESOURCES TABLE

| REAGENT or RESOURCE | SOURCE | IDENTIFIER |
| --- | --- | --- |
| **Antibodies** |  |  |
| Anti-Flag mAb | Sigma | F1804/clone M2 |
| anti-glyceraldehyde-3-phosphate dehydrogenase (GAPDH) PAb | AntGene | ANT325 |
| anti-β-tubulin mAb | Immuno Way | YM3030 |
| Alexa Fluor 488-conjugated goat anti-human IgG | Thermo Fisher Scientific | A11013 |
| Alexa Fluor 594-conjugated goat anti-mouse IgG | Thermo Fisher Scientific | A32742 |
| DyLight 594-conjugated goat anti-human IgG | Thermo Fisher Scientific | SA5-10136 |
| Horseradish peroxidase (HRP)-conjugated secondary antibody AffiniPure Goat Anti-Mouse IgG (H+L) | Jackson Immuno Research | 115-035-003 |
| Horseradish peroxidase (HRP)-conjugated secondary antibody AffiniPure Goat Anti-Rabbit IgG (H+L) | Jackson Immuno Research | 111-035-003 |
| **Bacterial and virus strains** |  |  |
| DH5α Chemically Competent Cell | TSINGKE | TSC-C14 |
| TransStbl3 Chemically Competent Cell | transgen | CD521-01 |
| G* $\Delta$ G-VSV(indicated as VSV-dG-GFP in this study) | Kerafast | Cat# EH1020-PM |
| VSV-dG-GFP-Fluc | This manuscript | N/A |
| **Chemicals, peptides, and recombinant proteins** |  |  |
| NeoCoV RBD (380-585aa) | This manuscript | N/A |
| PDF-2180 RBD (381-586aa) | This manuscript | N/A |
| SARS-CoV-2 RBD (331-524aa) | This manuscript | N/A |
| NL63 RBD (481-616aa) | This manuscript | N/A |
| *Rousettus aegyptiacus* ACE2 (18-740aa) | This manuscript | N/A |
| *Rhinolophus sinicus* ACE2 (18-740aa) | This manuscript | N/A |
| *Pipistrellus kuhlii* ACE2 (20-738aa) | This manuscript | N/A |
| UltraPure 1M Tris-HCl, pH 8.0 | Thermo Scientific | 15568025 |
| Hoechst 33342 | Sigma | 14533 |
| Trypsin from bovine pancreas (TPCK Treated) | Sigma | T8802 |
| Soybean trypsin inhibitor | Sigma-Aldrich | T6414 |
| Polybrene | Beyotime | C035 |
| PMSF | Beyotime | ST506 |
| Critical commercial assays |  |  |
| GeneTwin reagent | Biomed | TG101-01 |
| SMM 293-TII Expression Medium | Sino Biological | M293TII |
| Pierce Protein A/G Plus Agarose | Thermo Scientific | 20424 |
| Strep-Tactin XT 4Flow high-capacity resin | IBA | 2-5030-002 |
| Omni-Easy Instant BCA Protein Assay Kit | Epizyme | ZJ102 |
| Protein A (ProA) biosensors | ForteBio | 18-5010 |
| soybean trypsin inhibitor | Sigma-Aldrich | T6414 |
| Bright-Glo Luciferase Assay Kit | Promega | E2620 |
| Omni-ECL Femto Light Chemiluminescence Kit | EpiZyme | SQ201 |
| **Experimental models: Cell lines** |  |  |
| HEK293T | ATCC | CRL-3216 |
| Bat epithelial cell line Tb 1 Lu | ATCC | CCL-88 |
| I1-Hybridoma (CRL-2700) cell line | ATCC | CRL-2700 |
| Recombinant DNA |  |  |
| WT or mutated bats ACE2 orthologs | This manuscript | N/A |
| WT or mutated non-bat mammalian ACE2 orthologs | This manuscript | N/A |
| NeoCoV spike | This manuscript | AGY29650.2 |
| PDF-2180 spike | This manuscript | YP_009361857.1 |
| SARS-CoV-2 (D614G) | This manuscript | YP_009724390.1 |
| NL63 | This manuscript | APF29071.1 |
| **Software and algorithms** |  |  |
| Octet Data Analysis software | ForteBio | Version: 12.2.0.20 |
| IQ-TREE | J. Trifinopoulos et al., 2016 | http://igtree.cibiv.univie.ac.at/ |
| MEGA-X | Sudhir Kumar et al., 2018 | https://www.megasoftware.net/ |
| ClustalW | Kyoto University Bioinformatics Center | https://www.genome.jp/tools-bin/clustalw |
| Geneious Prime software | Geneious by Dotmatics | https://www.geneious.com/prime/ |
| iTOL |  | https://itol.embl.de/ |
| UCSF Chimera | Pettersen et al., 2004 | https://www.cgl.ucsf.edu/chimera |
| UCSF ChimeraX | Goddard et al., 2018 | https://www.rbvi.ucsf.edu/chimerax/ |
| FlowJo | FlowJo, LLC | https://www.flowjo.com/ |
| GraphPad Prism 8 | GraphPad Software | https://www.graphpad.com/scientific-software/prism/ |

## RESOURCE AVAILABILITY

### Lead contact

Further information and requests for resources and reagents should be directed to and will be fulfilled by the lead contact, Huan Yan

### Materials availability

All reagents generated in this study are available from the lead contact with a completed Materials Transfer Agreement.

### Data and code availability

- Supplemental Tables will be available from Mendeley Data.
- This study did not generate custom computer code.
- Any additional information required to reanalyze the data reported in this paper is available from the lead contact upon request.

## EXPERIMENTAL MODEL AND SUBJECT DETAILS

### Cell lines

HEK293T (CRL-3216) and the bat epithelial cell line Tb 1 Lu (CCL-88) were purchased from American Type Culture Collection (ATCC). Cells were maintained by Dulbecco’s Modified Eagle Medium (DMEM, Monad, China) with 1% PS (Penicillin/Streptomycin) and 10% Fetal Bovine Serum. An I1-Hybridoma (CRL-2700) cell line secreting a neutralizing mouse monoclonal antibody targeting the VSV glycoprotein (VSVG) was cultured in Minimum Essential Medium (MEM) with Earles’s balances salts and 2.0 mM of L-glutamine (Gibico) and 10% FBS. All the cells were cultured at 37°C with 5% CO2 with regular passage every 2-3 days.

### Plasmids

Plasmids expressing WT or mutated bats ACE2 orthologs were generated by inserting human codon-optimized sequences with/without specific mutations into a lentiviral transfer vector (pLVX-EF1a-Puro, Genewiz) with C-terminus 3×Flag (DYKDHD-G-DYKDHD-I-DYKDDDDK). Human codon-optimized sequences of all non-bat mammalian ACE2 and their mutants were cloned into a vector (pLVX-IRES-zsGreen) with a C-terminal Flag tag (DYKDDDDK) ^26^. Human codon-optimized spike sequences of NeoCoV (AGY29650.2), PDF-2180 (YP_009361857.1), SARS-CoV-2 (YP_009724390.1) carrying D614G mutation, and NL63 (APF29071.1) were cloned into the PCAGGS vector with C terminal deletions (13-15aa) to improve the pseudovirus assembly efficiency. The plasmids expressing the recombinant CoVs RBD-hFc fusion proteins were constructed by inserting NeoCoV RBD (380-585aa), PDF-2180 RBD (381-586aa), SARS-CoV-2 RBD (331-524aa) and NL63 RBD (481-616aa) coding sequences into the pCAGGS vector containing an N-terminal CD5 secretion signal peptide (MPMGSLQPLATLYLLGMLVASVL) and a C-terminal hFc tag for purification and detection. The plasmids expressing bats ACE2 ectodomain proteins were generated by inserting WT or mutated sequences of *Rousettus_aegyptiacus* (18-740aa), *Rhinolophus_sinicus* (18-740aa), *and Pipistrellus_kuhlii* (20-738aa) into the pCAGGS vector with an N-terminal CD5 secretion signal peptide and a C-terminal twin-strep-3×Flag tag (WSHPQFEKGGGSGGGSGGSAWSHPQFEKGGGRSDYKDHDGDYKDHDIDYKDDDDK).

## METHOD DETAILS

### Protein expression and purification

HEK293T cells were transfected with different protein-expressing plasmids through the GeneTwin reagent (Biomed, TG101-01). At 4-6 hours post-transfection, the medium of the transfected cells was replenished with the SMM 293-TII Expression Medium (Sino Biological, M293TII), and the protein-containing supernatant was harvested every three days for 2-3 batches. Recombinant RBD-hFc proteins were captured by Pierce Protein A/G Plus Agarose (Thermo Scientific, 20424), washed by wash buffer (100 mM Tris/HCl, pH 8.0, 150 mM NaCl, 1 mM EDTA), eluted with pH 3.0 Glycine buffer (100 mM in H2O), and then immediately balanced by 1/10 volume of UltraPure 1M Tris-HCI, pH 8.0 (15568025, Thermo Scientific). Proteins with twin-strep tag were captured by Strep-Tactin XT 4Flow high-capacity resin (IBA, 2-5030-002), washed by wash buffer, and then eluted by buffer BXT (100 mM Tris/HCl, pH 8.0, 150 mM NaCl, 1 mM EDTA, 50 mM biotin). The eluted proteins were concentrated by Ultrafiltration tubes, buffer changed with PBS, and stored at −80°C. Protein concentrations were determined by the Omni-Easy Instant BCA Protein Assay Kit (Epizyme, ZJ102).

### Coronavirus RBD-hFc live-cell binding assays

Different coronavirus RBD-hFc recombinant proteins were diluted in DMEM at indicated concentrations and incubated with HEK293T cells expressing different ACE2 for 30 mins at 37°C at 36 hours post-transfection. After binding, cells were washed once by Hanks’ Balanced Salt Solution (HBSS) and then incubated with 2 μg/mL of Alexa Fluor 488-conjugated goat anti-human IgG (Thermo Fisher Scientific; A11013) or DyLight 594-conjugated goat anti-human IgG (Thermo Fisher Scientific; SA5-10136) diluted in PBS/1% BSA for 1 hour at 37°C. Next, cells were washed once by HBSS and then incubated with Hoechst 33342 (1:10,000 dilution in HBSS) for 30 mins at 37°C to stain the nucleus. Images were captured by a fluorescence microscope (MI52-N). The relative fluorescence intensities (RFU) of the stained cells were determined by a Varioskan LUX Multi-well Luminometer (Thermo Scientific). For flow cytometry analysis, the stained cells were detached by 5 mM of EDTA/PBS and analyzed with a CytoFLEX Flow Cytometer (Beckman). 10,000 events in a gated live cell population (based on SSC/FSC) were analyzed for all samples. HEK293T cells transfected with empty vector plasmid were used as negative controls.

### Biolayer interferometry (BLI) binding assays

BLI assays were performed on the Octet RED96 instrument (Molecular Devices) following the manufacturer’s instructions. In general, RBD-hFc recombinant proteins (20 μg/mL) were immobilized on the Protein A (ProA) biosensors (ForteBio, 18-5010) and incubated with 2-fold serial-diluted bat ACE2-ectodomain proteins starting from 500 nM in the kinetic buffer (PBST). The background was set with a kinetic buffer without the ACE2-ectodomain proteins. The kinetic parameters and binding affinities between the RBD-hFc and different bat ACE2 were analyzed by Octet Data Analysis software 12.2.0.20 through curve-fitting kinetic or steady-state analysis.

### Pseudovirus production and titration

VSV-dG-based pseudoviruses carrying CoVs spike proteins were produced based on a modified protocol as previously described^53^. In general, HEK293T cells were transfected with CoVs spike protein-expressing plasmids. At 24 hours post-transfection, cells were transduced with 1.5×10^6^ TCID50 VSV-G glycoprotein-deficient VSV expressing GFP and firefly luciferase (VSV-dG-GFP-fLuc, generated in our lab) diluted in DMEM with 8 μg/mL polybrene for 4-6 hours at 37°C. After three times of PBS wash, the culture medium was replenished with DMEM+10% FBS or SMM 293-TII Expression Medium (Sino Biological, M293TII) containing VSV neutralizing antibody (from I1-mouse hybridoma). Twenty-four hours later, the virus-containing supernatant was clarified through centrifugation at 4,000 rpm for 5 mins at 4°C and then stored at −80°C. TCID50 of pseudotyped viruses were determined based on three-fold serial dilution-based infection assays on HEK293T-bat40ACE2 cells for NeoCoV and PDF-2180 S pseudotypes, and 293T-hACE2 cells for NL63 and SARS-CoV-2 S pseudotypes. TCID50 was calculated according to the Reed-Muench method^54,55^.

### Pseudovirus entry assay

Pseudovirus entry assays were conducted on HEK293T or Tb 1 Lu cells transiently expressing WT or mutant ACE2 orthologs at 36 hours post-transfection. In general, 3×10^4^ trypsinized cells were incubated with pseudovirus (1.5×10^5^ TCID50/100 μL) in a 96-well plate to allow attachment and viral entry. TPCK-trypsin (Sigma-Aldrich, T8802) treatment was conducted before NeoCoV and PDF-2180 pseudovirus entry assay. In this case, pseudovirus produced in Serum-free SMM 293-TII Expression Medium were incubated with 100 μg/mL TPCK-treated trypsin for 10 mins at room temperature, followed by neutralization with 100 μg/mL soybean trypsin inhibitor (Sigma-Aldrich, T6414). The intracellular luciferase activity was measured by Bright-Glo Luciferase Assay Kit (Promega, E2620) and detected with a GloMax 20/20 Luminometer (Promega) at 18 hours post-infection. GFP intensity was analyzed by a fluorescence microscope (Mshot, MI52-N).

### Western blot

For Western blot analysis, cells were lysed with 1% TritonX/PBS+1 mM PMSF (Beyotime, ST506) for 10 mins at 4°C, then clarified through centrifugation of 12000 rpm for 5 mins at 4°C. The clarified cell lysate was mixed with the 1/5 volume of 5×SDS loading buffer and incubated at 98°C for 10 mins. After gel electrophoresis and membrane transfer, the PVDF-membrane blots were blocked with 5% skimmed milk in PBST for 2 hours at room temperature and then incubated 1 μg/mL anti-Flag mAb (Sigma, F1804), anti-glyceraldehyde-3-phosphate dehydrogenase (GAPDH) (AntGene, ANT325) PAb or anti-β-tubulin (Immmuno Way, YM3030) mAb diluted in PBST containing 1% milk overnight at 4°C. After three times washing with PBST, the blots were incubated with Horseradish peroxidase (HRP)-conjugated secondary antibody AffiniPure Goat Anti-Mouse or Rabbit IgG (H+L) (Jackson Immuno Research, 115-035-003 or 111-035-003) in 1% skim milk in PBST and incubated for one hour at room temperature. The blots were then washed three times by PBST and then visualized using an Omni-ECL Femto Light Chemiluminescence Kit (EpiZyme, SQ201) by a ChemiDoc MP Imaging System (Bio-Rad).

### Immunofluorescence assay

Immunofluorescence assays were conducted to verify the expression levels of ACE2 with C-terminal fused 3×Flag. In general, the transfected cells were incubated with 100% methanol for 10 mins at room temperature for fixation and permeabilization. Cells were then incubated with a mouse antibody M2 (Sigma-Aldrich, F1804) diluted in PBS/1% BSA for one hour at 37°C, followed by extensive wash and the incubation of secondary antibody of Alexa Fluor 594-conjugated goat anti-mouse IgG (Thermo Fisher Scientific, A32742) diluted in 1% BSA/PBS for one hour at 37°C. Before visualization, the nucleus was stained blue with Hoechst 33342 reagent (1:5,000 dilution in PBS). Images were captured and merged with a fluorescence microscope (Mshot, MI52-N).

### Bioinformatic and structural analysis

Sequence alignments of different bats ACE2 or non-bat mammalian ACE2 were performed either by the MUSCLE algorithm by MEGA-X (version 10.1.8) or ClustalW (https://www.genome.jp/tools-bin/clustalw) software. The residue usage frequency (sequence logo) and mean hydrophobicity of all ACE2 sequences were generated by the Geneious Prime software. Phylogenetic trees were produced using the maximal likelihood method in IQ-TREE (http://igtree.cibiv.univie.ac.at/) (1000 Bootstraps) and polished with iTOL (v6) (https://itol.embl.de/)^56^. The structural were shown by ChimeraX based on SARS-CoV-2 RBD & human ACE2 (PDB: 6M0J), NL63 RBD & human ACE2 (PDB: 3KBH), NeoCoV RBD & Ppip ACE2 (PDB: 7WPO) and PDF-2180 RBD & Ppip ACE2 (PDB: 7WPZ). RBD binding footprints and interaction details were analyzed and demonstrated using the UCSF ChimeraX^57^. Structural representatives of NeoCoV RBD interacting with WT or mutated Ppip ACE2 were analyzed using the UCSF Chimera X. The colored sequence conservation of 102 mammals ACE2 was demonstrated by the UCSF Chimera based on multi-sequence alignments data generated by MEGA-X.

## QUANTIFICATION AND STATISTICAL ANALYSIS

### Statistical Analysis

Most experiments were conducted 2-3 times with 3 or 4 biological repeats. Representative results were shown. Data were presented by MEAN±SD or MEAN±SEM as indicated in the figure legends. Unpaired two-tailed t-tests were conducted for all statistical analyses using GraphPad Prism 8. *P*<0.05 was considered significant. \**p*<0.05, \*\**p* <0.01, \*\*\**p* <0.005, and **** *p* <0.001.

---

**Figure S1.**
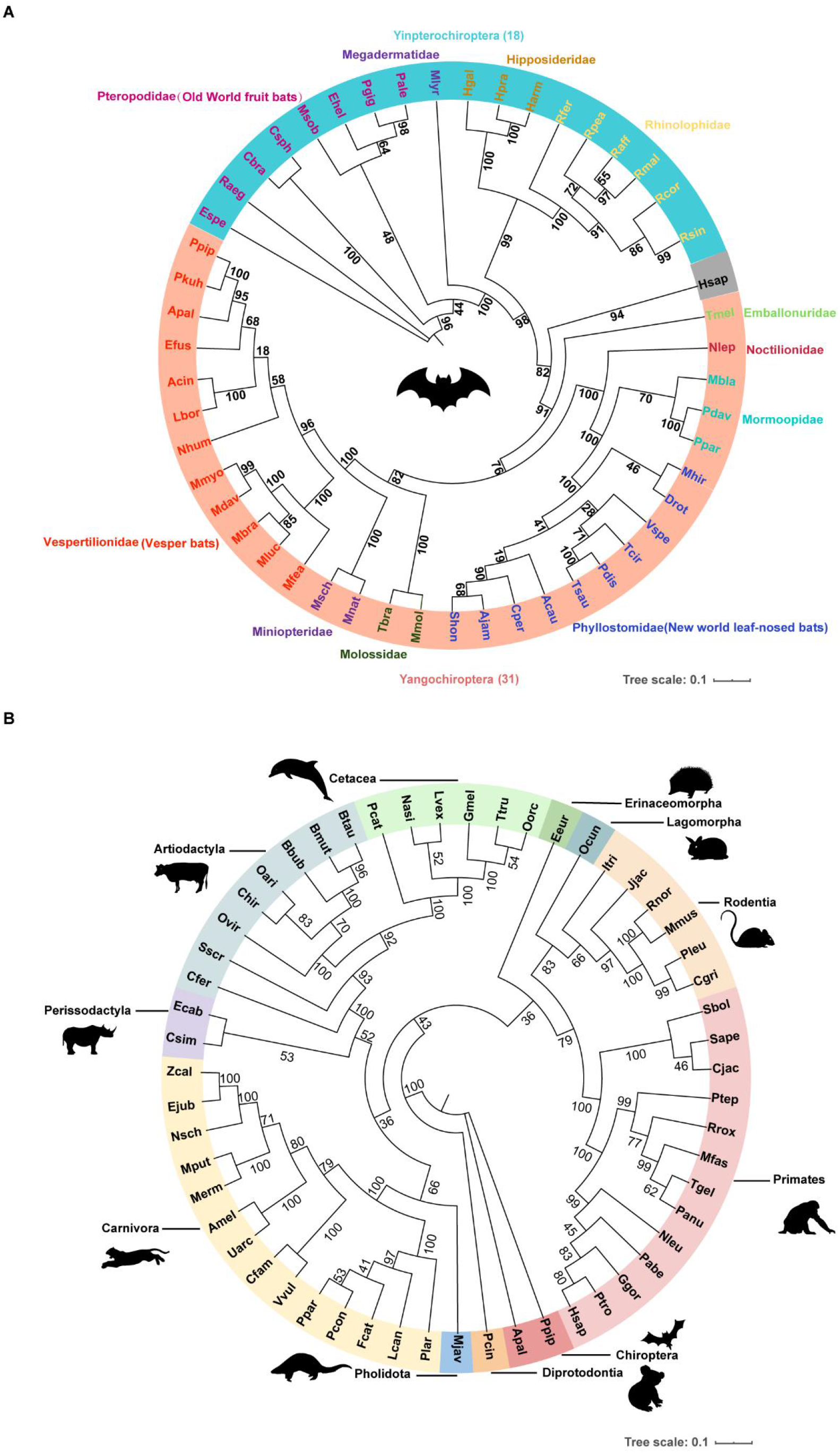
The phylogenetic trees of the bats and mammalian ACE2 tested in this study. (**A-B**) The phylogenetic trees were generated by the IQ-TREE based on the ACE2 protein sequences from the 49 bats species (**A**) or 55 mammals (including two bats) (**B**). Species of the same order or suborder were highlighted with different background colors. The GenBank accession numbers and protein sequences were summarized in **Supplementary Table 1**.

**Figure S2.**
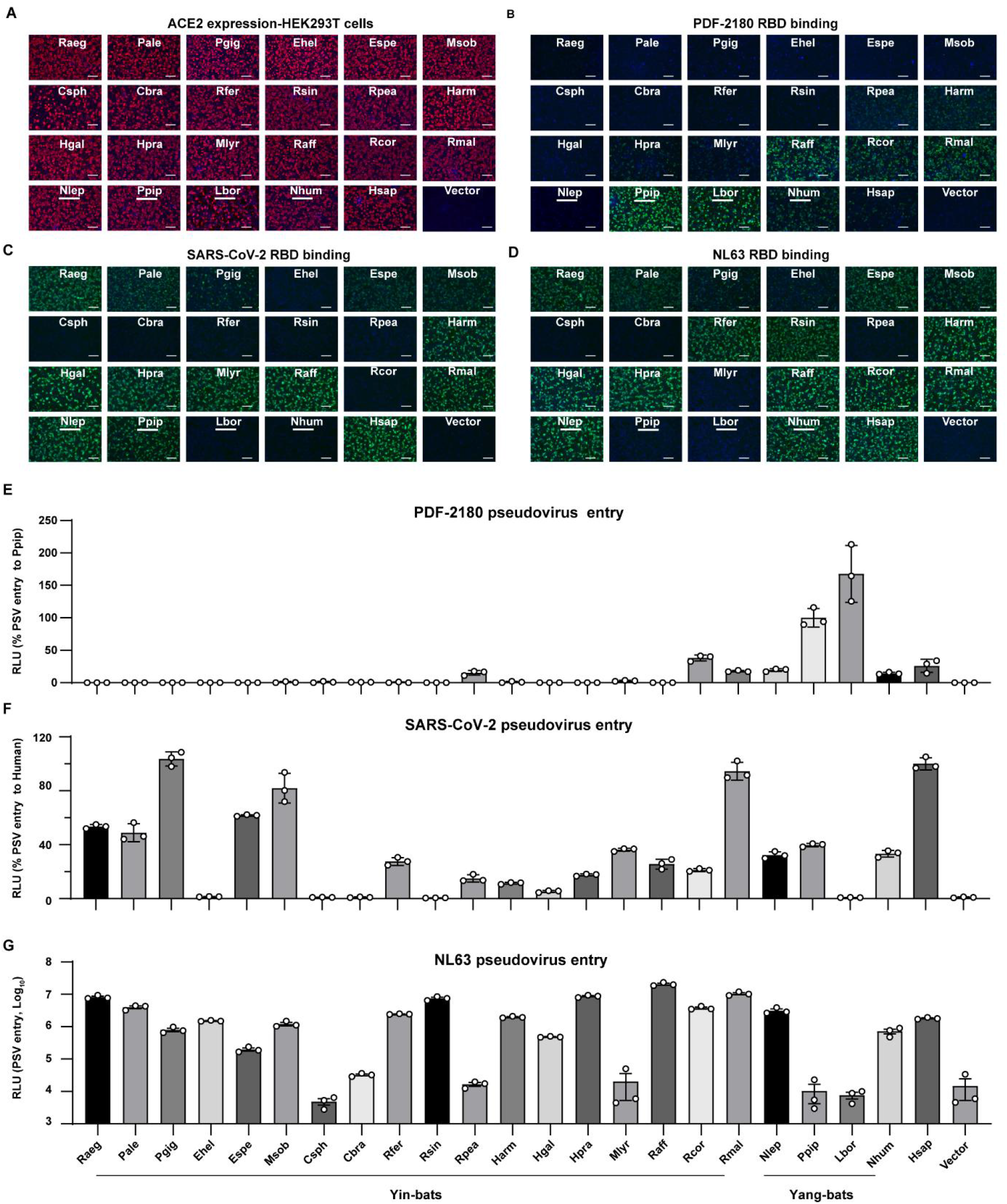
The RBD binding and pseudoviruses entry efficiency of ACE2-using CoVs in HEK293T cells transiently expressing the indicated bats ACE2 orthologs. (**A**) The expression level of ACE2 orthologs examined by immunofluorescence assay detecting the C terminal fused 3×Flag. (**B-G**) The RBD binding (**B-D**) and pseudoviruses entry efficiency (**E-G**) of PDF-2180 (**B** and **E**), SARS-CoV-2 (**C** and **F**), NL63 (**D** and **G**) in HEK293T cells transiently expressing the indicated bats ACE2. Data are presented as mean ± SD for n=3 biologically independent cells for E and F. Data are presented as mean ± SEM for n=3 biologically independent cells for G. Data representative of two independent experiments. RLU: relative luciferase unit. Scale bar represents 100 μm for **A**, **B**, **C**, and **D**.

**Figure S3.**
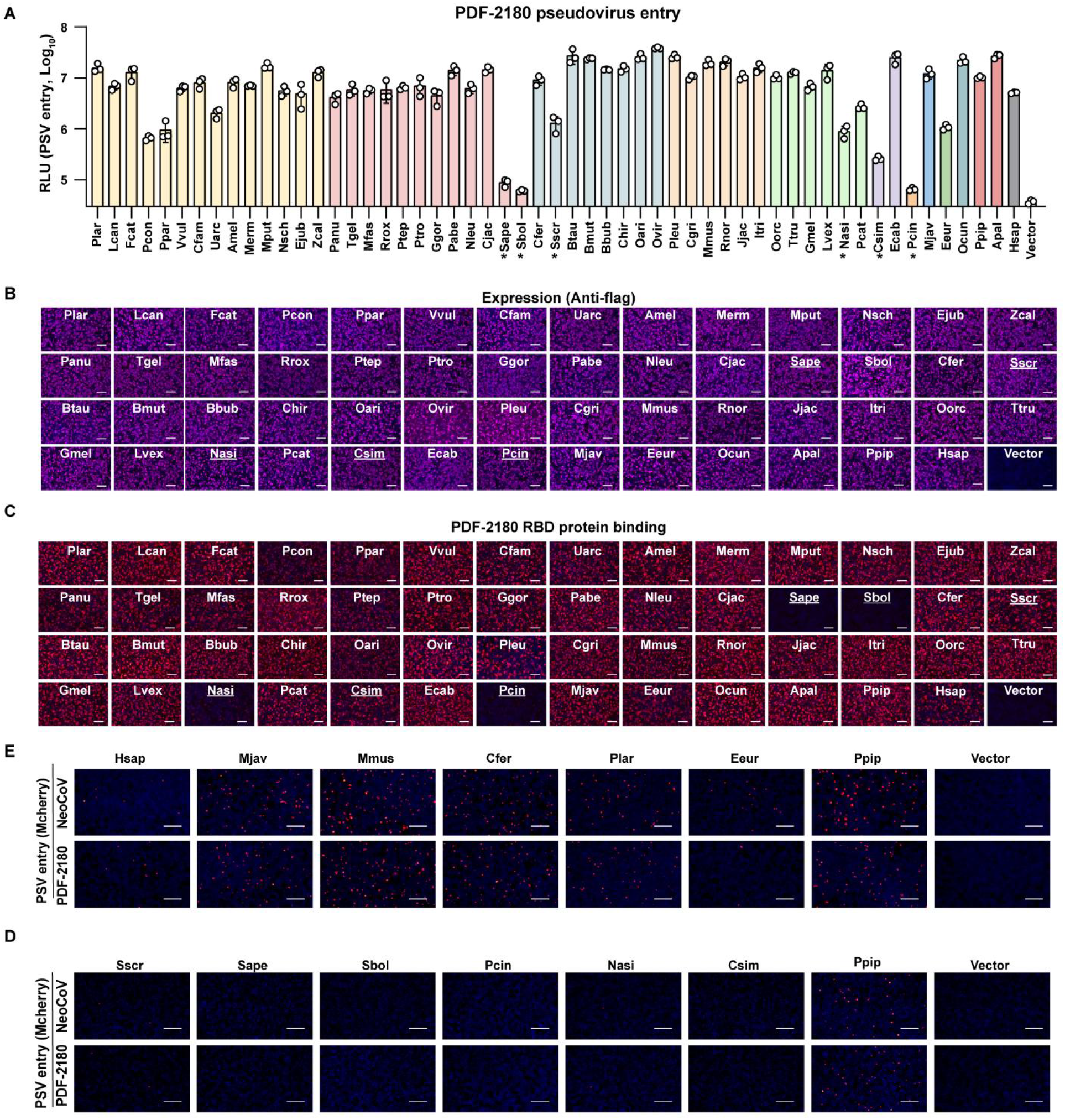
NeoCoV and PDF-2180 can recognize most ACE2 orthologs from non-bat mammals. (**A**) Entry efficiency of PDF-2180 pseudovirus in HEK293T cells transiently expressing mammalian ACE2. *: RLU<20% RLU_Ppip_ of NeoCoV pseudovirus entry. Species belonging to different orders were indicated with different background colors. (**B**) The expression level of ACE2 orthologs in HEK293T cells as indicated by immunofluorescence assay detecting the C-terminal Flag tag. (**C**) PDF-2180 RBD-hFc binding efficiency in HEK293T cells expressing indicated ACE2 from non-bat mammals as indicated by immunofluorescence assay detecting the hFc. (**D-E**) Entry efficiency of NeoCoV and PDF-2180 pseudoviruses (mcherry) in HEK293T cells transiently expressing indicated ACE2 orthologs from CoV host-related species (**D**) or non-permissive species (**E**). Data are presented as mean ± SD for n=3 biologically independent cells. Data representative of two independent experiments. RLU: relative luciferase unit. Scale bar represents 100 μm for **B-C** and 200 μm for **D-E**.

**Figure S4.**
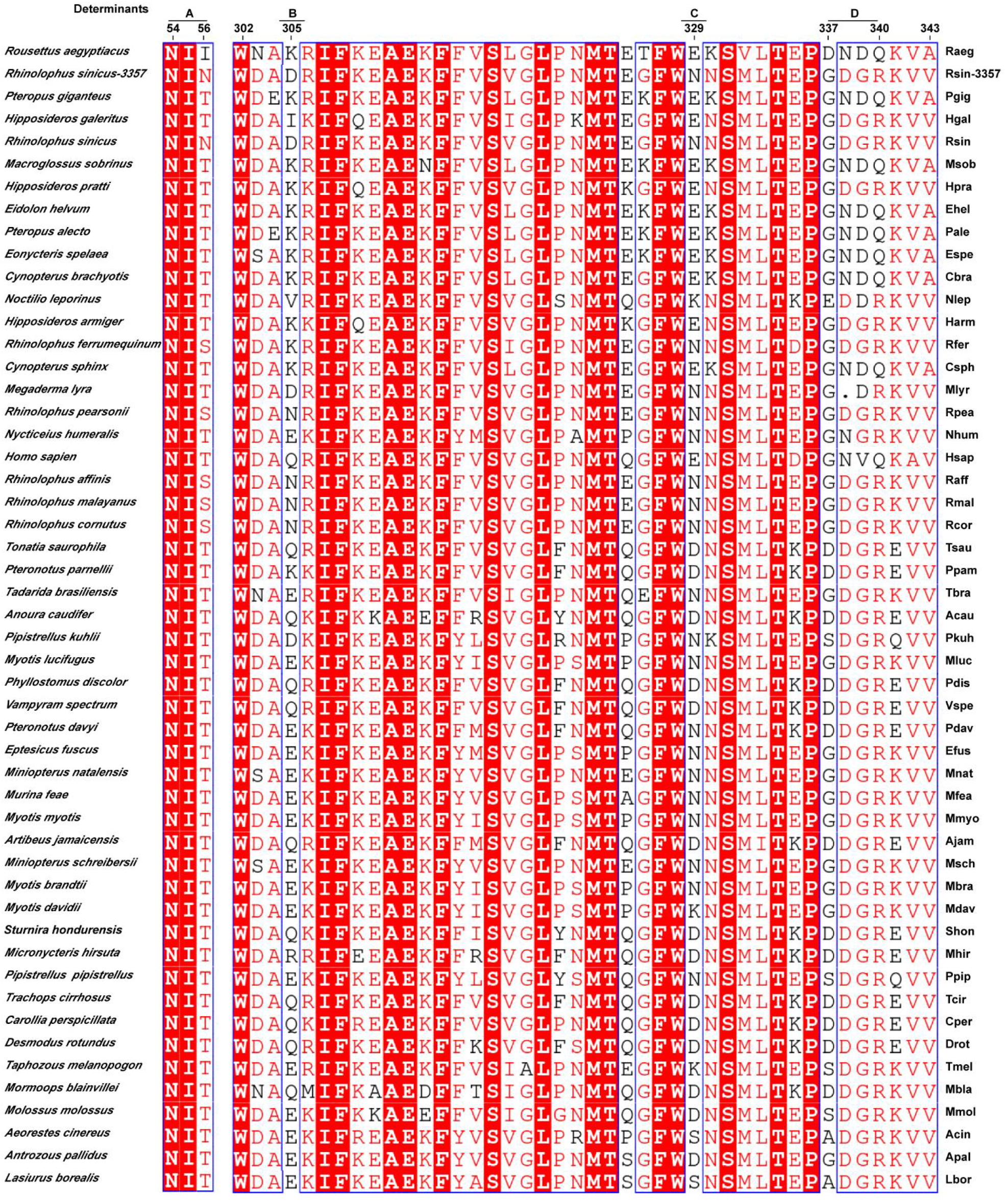
The multi-sequence alignments of viral binding loops of bat ACE2 orthologs. The sequence alignment analysis was conducted based on viral binding loops of ACE2 by the Clustal W and rendered with ESPript. Identical residues are highlighted with a red background, and similar residues are colored red in blue boxes.

**Figure S5.**
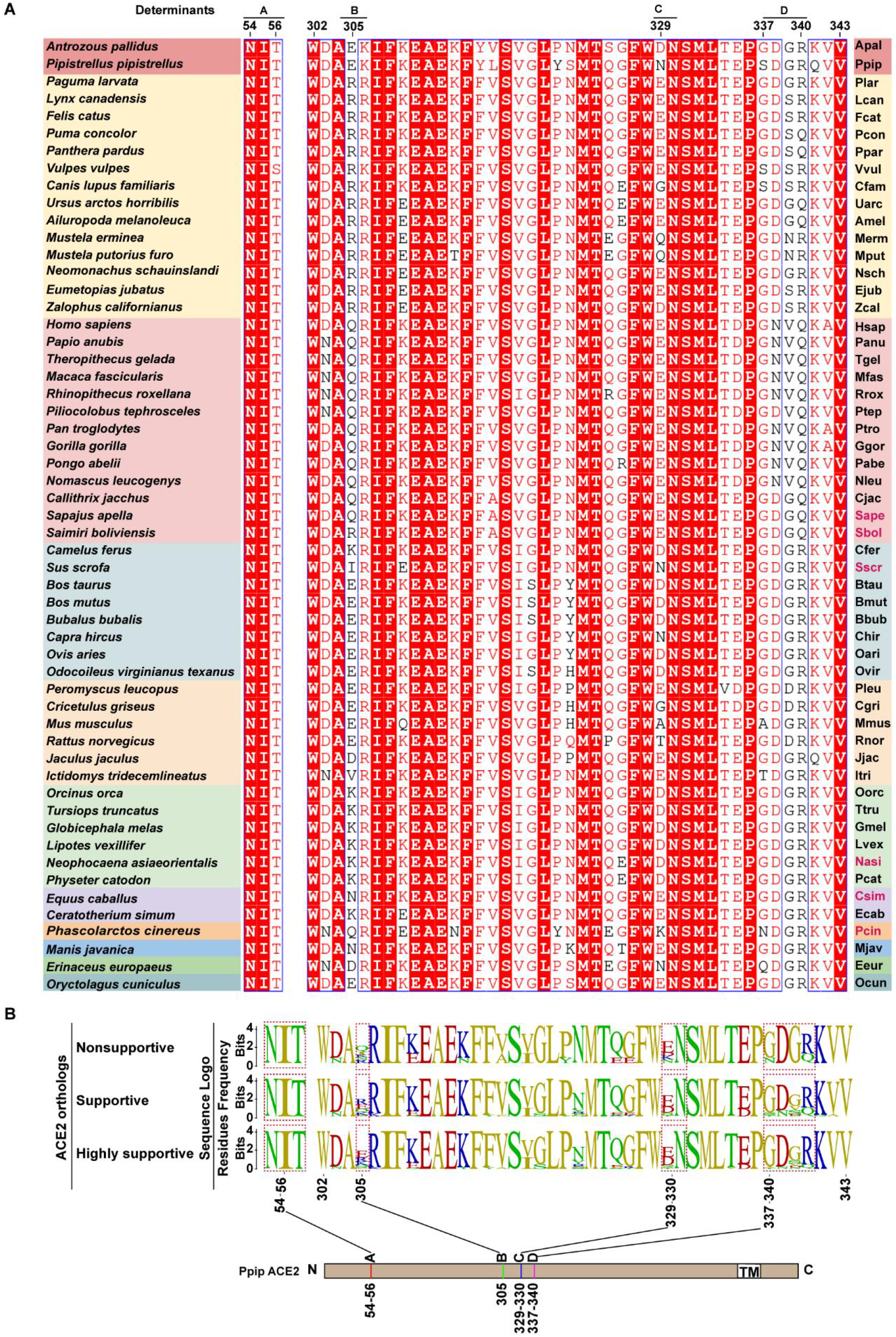
The multi-sequence alignments and sequence conservation analysis of viral binding loops of ACE2 orthologs from 55 mammals. (**A**) Multi-sequence alignment based on viral binding loops from 55 mammals (including two bats) by the Clustal W and rendered with ESPript. Identical residues were highlighted in red, and similar residues were in the blue frames. (**B**) Sequence conservation analysis of the ACE2 orthologs grouped by their ability to support NeoCoV entry. Upper: non-supportive (<20% RLU_Ppip_, 6 species); middle: supportive (> 20% RLU_Ppip_, 49 species); lower: highly supportive (>100% RLU_Ppip_, 30 species).

**Figure S6.**
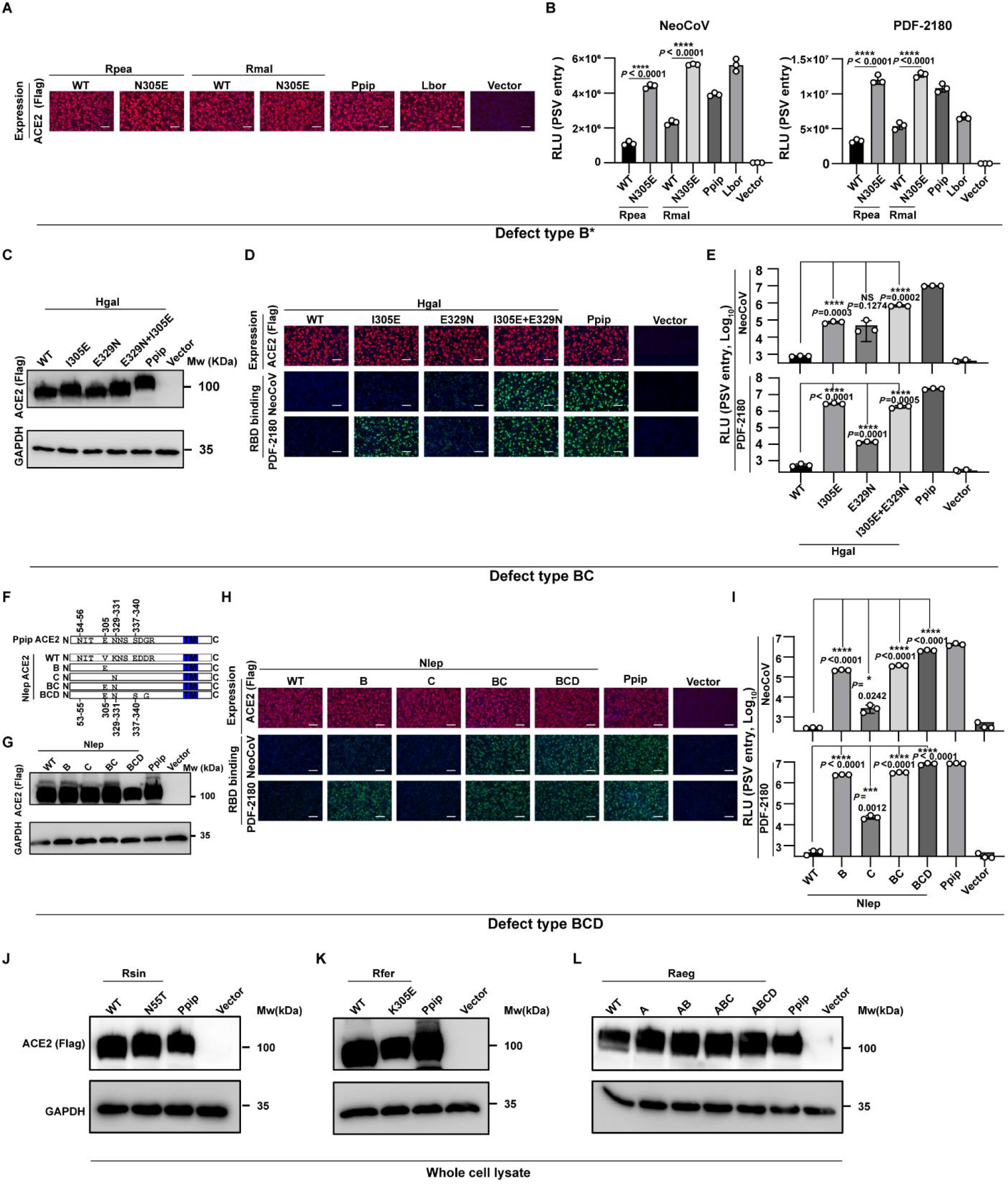
Verification of bats ACE2 mutants with improved functionality to support NeoCoV/PDF-2180 RBD binding and pseudoviruses entry. (**A**) Expression levels of the WT and mutated Rpea and Rmal ACE2 by immunofluorescence in HEK293T cells. (**B**) The NeoCoV and PDF-2180 pseudoviruses entry efficiency supported by WT and mutated Rpea and Rmal ACE2. (**C**) Western blot analysis of the WT and mutated Hgal ACE2 expression in HEK293T cells. (**D**) Expression levels and RBD binding supporting ability of WT and mutated Hgal ACE2. (**E**) The NeoCoV and PDF-2180 pseudoviruses entry efficiency supported by WT and mutated Hgal ACE2. (**F**) Schematic illustration of Nlep ACE2 swap mutants carrying the indicated PpipACE2 counterparts. (**G**) Western blot analysis of the expression levels of WT and mutated Nlep ACE2 in HEK293T cells. (**H**) The binding efficiency of NeoCoV and PDF-2180 RBD supported by WT and mutated Nlep ACE2. (**I**) Entry efficiency of NeoCoV and PDF-2180 pseudoviruses supported by WT and mutated Nlep ACE2. (**J-L**) Western blot analysis of the expression levels of WT and mutated Rsin (**J**), Rfer (**K**), and Raeg (**L**) ACE2 in HEK293T cells. Data are presented as mean ± SD for n=3 biologically independent cells for B, E, I. Data representative of three independent experiments for A-E and G-I. Data representative of two experiments for J-L. Two-tailed unpaired Student’ s t-test; * p<0.05, ** p <0.01, *** p <0.005, and **** p <0.001. NS: not significant. RLU: relative luciferase unit. Mw: molecular weight. Scale bar represents 100 μm for **A**,**D**, and **H**.

**Figure S7.**
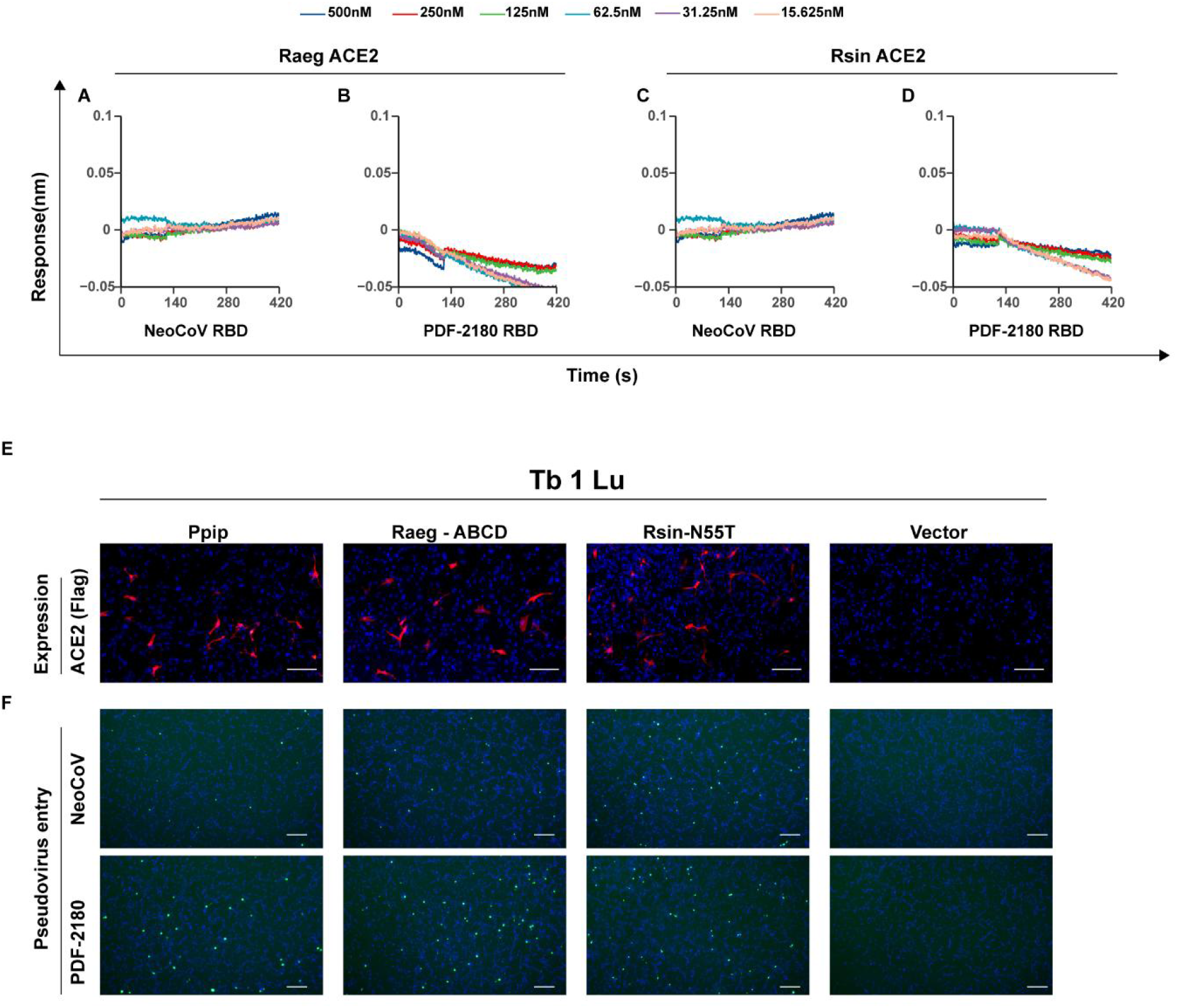
Verification of the receptor function of Raeg and Rsin ACE2 mutants with improved receptor recognition. (**A-D**) BLI assays analyzing the binding kinetics between NeoCoV-RBD-hFc/PDF-2180-RBD-hFc and WT Raeg and Rsin ACE2 ectodomain proteins. (**E-F**) Verification of the gain of receptor function of Raeg and Rsin ACE2 mutants in Tb 1 Lu bat cell line. Immunofluorescence analysis of ACE2 expression level of Ppip and mutated Raeg and Rsin ACE2 in Tb1 Lu cells (**E**). NeoCoV and PDF-2180 pseudoviruses entry efficiency at 16 h post-infection as indicated by the GFP intensity (**F**). Data representative of two independent experiments. Scale bar represents 100 μm for **E** and 300μm for **F**.

**Figure S8.**
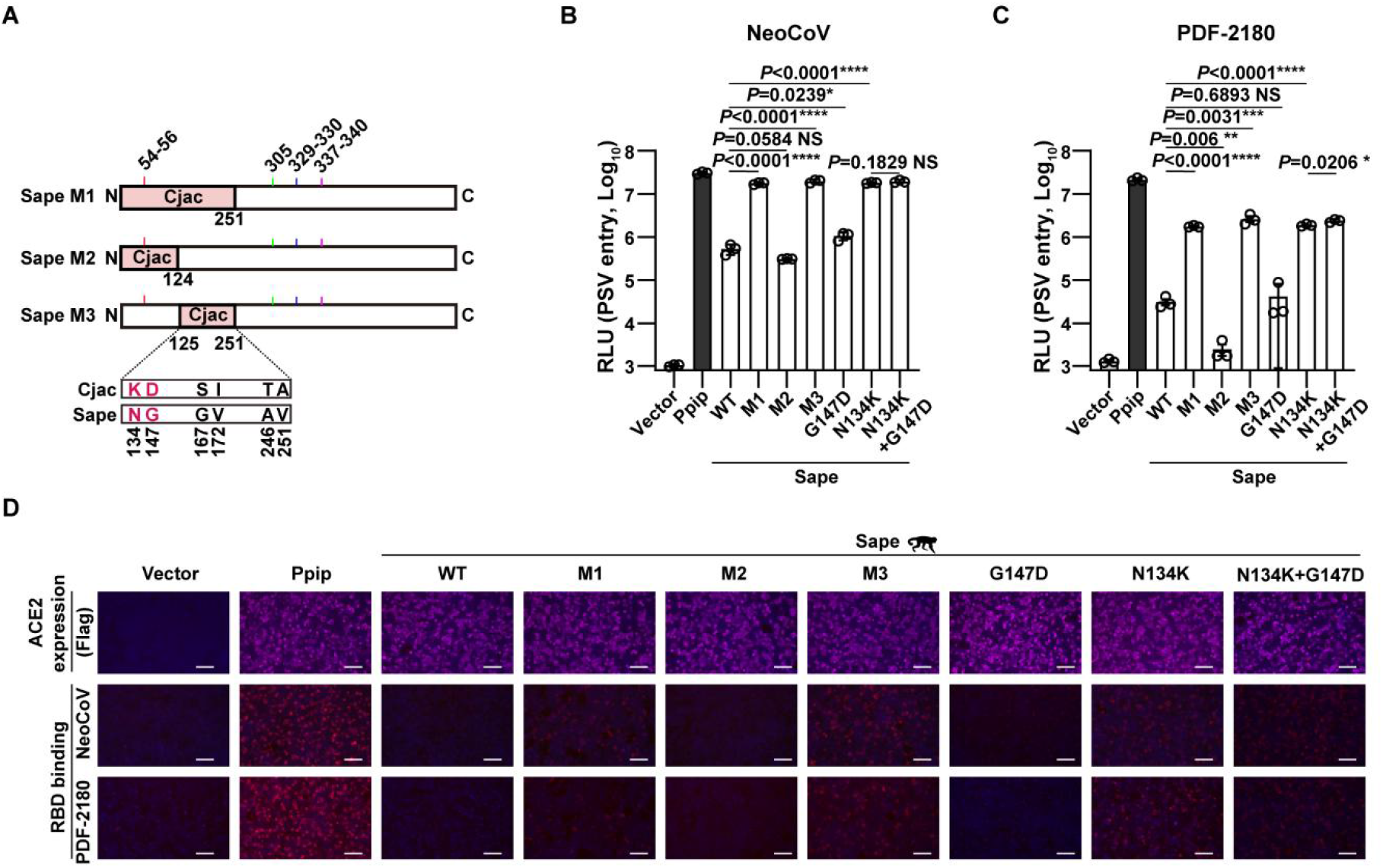
Residue K134 is a critical determinant restricting Sape ACE2 from supporting NeoCoV and PDF-2180 RBD binding and pseudoviruses entry. (**A**) Schematic illustration of Sape ACE2 swap mutants carrying the indicated Cjac ACE2 counterparts. (**B**-**D**) Identification of the critical determinants restricting Sape ACE2 from supporting NeoCoV/PDF-2180 **pseudoviruses** entry (**B-C**) and RBD binding (**D**) in HEK293T cells. Data are presented as mean ±SD for n=3 biologically independent cells for **B** and **C**. Data representative of two independent experiments. Two-tailed unpaired Student’s t-test; * p<0.05, ** p <0.01, *** p <0.005, and **** p <0.001. NS: not significant. RLU: relative luciferase unit. Scale bar represents 100 μm for **D**.

## Reference

1 Zaki, A. M., van Boheemen, S., Bestebroer, T. M., Osterhaus, A. D. & Fouchier, R. A. Isolation of a novel coronavirus from a man with pneumonia in Saudi Arabia. N Engl J Med 367, 1814–1820, doi:10.1056/NEJMoa1211721 (2012).

2 Zhou, P. et al. A pneumonia outbreak associated with a new coronavirus of probable bat origin. Nature 579, 270–273, doi:10.1038/s41586-020-2012-7 (2020).

3 Chen, L. et al. RNA based mNGS approach identifies a novel human coronavirus from two individual pneumonia cases in 2019 Wuhan outbreak. Emerg Microbes Infect 9, 313–319, doi:10.1080/22221751.2020.1725399 (2020).

4 Ksiazek, T. G. et al. A novel coronavirus associated with severe acute respiratory syndrome. N Engl J Med 348, 1953–1966, doi:10.1056/NEJMoa030781 (2003).

5 Ge, X. Y. et al. Isolation and characterization of a bat SARS-like coronavirus that uses the ACE2 receptor. Nature 503, 535–538, doi:10.1038/nature12711 (2013).

6 Latinne, A. et al. Origin and cross-species transmission of bat coronaviruses in China. Nat Commun 11, 4235, doi:10.1038/s41467-020-17687-3 (2020).

7 Li, W. et al. Bats are natural reservoirs of SARS-like coronaviruses. Science 310, 676–679, doi:10.1126/science.1118391 (2005).

8 Temmam, S. et al. Bat coronaviruses related to SARS-CoV-2 and infectious for human cells. Nature 604, 330–336, doi:10.1038/s41586-022-04532-4 (2022).

9 Wang, Q. et al. Bat origins of MERS-CoV supported by bat coronavirus HKU4 usage of human receptor CD26. Cell Host Microbe 16, 328–337, doi:10.1016/j.chom.2014.08.009 (2014).

10 Wong, A. C. P., Li, X., Lau, S. K. P. & Woo, P. C. Y. Global Epidemiology of Bat Coronaviruses. Viruses 11, doi:10.3390/v11020174 (2019).

11 WHO. MERS situation update, <https://www.emro.who.int/health-topics/mers-cov/mers-outbreaks.html> (2022).

12 Anthony, S. J. et al. Further Evidence for Bats as the Evolutionary Source of Middle East Respiratory Syndrome Coronavirus. mBio 8, doi:10.1128/mBio.00373-17 (2017).

13 Corman, V. M. et al. Rooting the phylogenetic tree of middle East respiratory syndrome coronavirus by characterization of a conspecific virus from an African bat. J Virol 88, 11297–11303, doi:10.1128/JVI.01498-14 (2014).

14 Geldenhuys, M. et al. A metagenomic viral discovery approach identifies potential zoonotic and novel mammalian viruses in Neoromicia bats within South Africa. PLoS One 13, e0194527, doi:10.1371/journal.pone.0194527 (2018).

15 Xiong, Q. et al. Close relatives of MERS-CoV in bats use ACE2 as their functional receptors. Nature, doi:10.1038/s41586-022-05513-3 (2022).

16 Li, W. et al. Angiotensin-converting enzyme 2 is a functional receptor for the SARS coronavirus. Nature 426, 450–454, doi:10.1038/nature02145 (2003).

17 Liu, K. et al. Binding and molecular basis of the bat coronavirus RaTG13 virus to ACE2 in humans and other species. Cell 184, 3438–3451 e3410, doi:10.1016/j.cell.2021.05.031 (2021).

18 Hofmann, H. et al. Human coronavirus NL63 uses the severe acute respiratory syndrome coronavirus receptor for cellular entry. Proc Natl Acad Sci USA 102, 7988–7993, doi:10.1073/pnas.0409465102 (2005).

19 Li, F. Receptor recognition mechanisms of coronaviruses: a decade of structural studies. J Virol 89, 1954–1964, doi:10.1128/JVI.02615-14 (2015).

20 Corman, V. M. et al. Characterization of a novel betacoronavirus related to middle East respiratory syndrome coronavirus in European hedgehogs. J Virol 88, 717–724, doi:10.1128/JVI.01600-13 (2014).

21 Ithete, N. L. et al. Close relative of human Middle East respiratory syndrome coronavirus in bat, South Africa. Emerg Infect Dis 19, 1697–1699, doi:10.3201/eid1910.130946 (2013).

22 Lau, S. K. P. et al. Identification of a Novel Betacoronavirus (Merbecovirus) in Amur Hedgehogs from China. Viruses 11, doi:10.3390/v11110980 (2019).

23 Douam, F. et al. Genetic Dissection of the Host Tropism of Human-Tropic Pathogens. Annu Rev Genet 49, 21–45, doi:10.1146/annurev-genet-112414-054823 (2015).

24 Menachery, V. D. et al. SARS-like WIV1-CoV poised for human emergence. Proc Natl Acad Sci U S A 113, 3048–3053, doi:10.1073/pnas.1517719113 (2016).

25 Wei, C. et al. Evidence for a mouse origin of the SARS-CoV-2 Omicron variant. J Genet Genomics 48, 1111–1121, doi:10.1016/j.jgg.2021.12.003 (2021).

26 Liu, Y. et al. Functional and genetic analysis of viral receptor ACE2 orthologs reveals a broad potential host range of SARS-CoV-2. Proc Natl Acad Sci U S A 118, doi:10.1073/pnas.2025373118 (2021).

27 King, S. B. & Singh, M. Comparative genomic analysis reveals varying levels of mammalian adaptation to coronavirus infections. PLoS Comput Biol 17, e1009560, doi:10.1371/journal.pcbi.1009560 (2021).

28 Letko, M. et al. Adaptive Evolution of MERS-CoV to Species Variation in DPP4. Cell Rep 24, 1730–1737, doi:10.1016/j.celrep.2018.07.045 (2018).

29 Liu, K. et al. Cross-species recognition of SARS-CoV-2 to bat ACE2. Proc Natl Acad Sci USA 118, doi:10.1073/pnas.2020216118 (2021).

30 Yan, H. et al. ACE2 receptor usage reveals variation in susceptibility to SARS-CoV and SARS-CoV-2 infection among bat species. Nat Ecol Evol 5, 600–608, doi:10.1038/s41559-021-01407-1 (2021).

31 Damas, J. et al. Broad host range of SARS-CoV-2 predicted by comparative and structural analysis of ACE2 in vertebrates. Proc Natl Acad Sci U S A 117, 22311–22322, doi:10.1073/pnas.2010146117 (2020).

32 Ren, W. et al. Comparative analysis reveals the species-specific genetic determinants of ACE2 required for SARS-CoV-2 entry. PLoS Pathog 17, e1009392, doi:10.1371/journal.ppat.1009392 (2021).

33 Zhao, X. et al. Broad and Differential Animal Angiotensin-Converting Enzyme 2 Receptor Usage by SARS-CoV-2. J Virol 94, doi:10.1128/JVI.00940-20 (2020).

34 Kistler, K. E., Huddleston, J. & Bedford, T. Rapid and parallel adaptive mutations in spike S1 drive clade success in SARS-CoV-2. Cell Host Microbe 30, 545–555 e544, doi:10.1016/j.chom.2022.03.018 (2022).

35 Oude Munnink, B. B. et al. Transmission of SARS-CoV-2 on mink farms between humans and mink and back to humans. Science 371, 172–177, doi:10.1126/science.abe5901 (2021).

36 Pickering, B. et al. Divergent SARS-CoV-2 variant emerges in white-tailed deer with deer-to-human transmission. Nat Microbiol, doi:10.1038/s41564-022-01268-9 (2022).

37 Huynh, J. et al. Evidence supporting a zoonotic origin of human coronavirus strain NL63. J Virol 86, 12816–12825, doi:10.1128/JVI.00906-12 (2012).

38 Haagmans, B. L. et al. Middle East respiratory syndrome coronavirus in dromedary camels: an outbreak investigation. Lancet Infect Dis 14, 140–145, doi:10.1016/S1473-3099(13)70690-X (2014).

39 Lam, T. T. et al. Identifying SARS-CoV-2-related coronaviruses in Malayan pangolins. Nature 583, 282–285, doi:10.1038/s41586-020-2169-0 (2020).

40 Song, H. D. et al. Cross-host evolution of severe acute respiratory syndrome coronavirus in palm civet and human. Proc Natl Acad Sci U S A 102, 2430–2435, doi:10.1073/pnas.0409608102 (2005).

41 Zhu, Z. et al. From SARS and MERS to COVID-19: a brief summary and comparison of severe acute respiratory infections caused by three highly pathogenic human coronaviruses. Respir Res 21, 224, doi:10.1186/s12931-020-01479-w (2020).

42 Breban, R., Riou, J. & Fontanet, A. Interhuman transmissibility of Middle East respiratory syndrome coronavirus: estimation of pandemic risk. Lancet 382, 694–699, doi:10.1016/S0140-6736(13)61492-0 (2013).

43 Lau, S. K. et al. Genetic characterization of Betacoronavirus lineage C viruses in bats reveals marked sequence divergence in the spike protein of pipistrellus bat coronavirus HKU5 in Japanese pipistrelle: implications for the origin of the novel Middle East respiratory syndrome coronavirus. J Virol 87, 8638–8650, doi:10.1128/JVI.01055-13 (2013).

44 Memish, Z. A. et al. Middle East respiratory syndrome coronavirus in bats, Saudi Arabia. Emerg Infect Dis 19, 1819–1823, doi:10.3201/eid1911.131172 (2013).

45 Mohd, H. A., Al-Tawfiq, J. A. & Memish, Z. A. Middle East Respiratory Syndrome Coronavirus (MERS-CoV) origin and animal reservoir. Virol J 13, 87, doi:10.1186/s12985-016-0544-0 (2016).

46 Zhang, Z., Shen, L. & Gu, X. Evolutionary Dynamics of MERS-CoV: Potential Recombination, Positive Selection and Transmission. Sci Rep 6, 25049, doi:10.1038/srep25049 (2016).

47 Fan, Y. et al. SARS-CoV-2 Omicron variant: recent progress and future perspectives. Signal Transduct Target Ther 7, 141, doi:10.1038/s41392-022-00997-x (2022).

48 Hu, B. et al. Discovery of a rich gene pool of bat SARS-related coronaviruses provides new insights into the origin of SARS coronavirus. PLoS Pathog 13, e1006698, doi:10.1371/journal.ppat.1006698 (2017).

49 Mehdipour, A. R. & Hummer, G. Dual nature of human ACE2 glycosylation in binding to SARS-CoV-2 spike. Proc Natl Acad Sci U S A 118, doi:10.1073/pnas.2100425118 (2021).

50 Tortorici, M. A. et al. Structure, receptor recognition, and antigenicity of the human coronavirus CCoV-HuPn-2018 spike glycoprotein. Cell 185, 2279–2291 e2217, doi:10.1016/j.cell.2022.05.019 (2022).

51 Wu, K., Peng, G., Wilken, M., Geraghty, R. J. & Li, F. Mechanisms of host receptor adaptation by severe acute respiratory syndrome coronavirus. J Biol Chem 287, 8904–8911, doi:10.1074/jbc.M111.325803 (2012).

52 Nomaguchi, M., Fujita, M., Miyazaki, Y. & Adachi, A. Viral tropism. Front Microbiol 3, 281, doi:10.3389/fmicb.2012.00281 (2012).

53 Whitt, M. A. Generation of VSV pseudotypes using recombinant DeltaG-VSV for studies on virus entry, identification of entry inhibitors, and immune responses to vaccines. J Virol Methods 169, 365–374, doi:10.1016/j.jviromet.2010.08.006 (2010).

54 Biacchesi, S. et al. Rapid human metapneumovirus microneutralization assay based on green fluorescent protein expression. J Virol Methods 128, 192–197, doi:10.1016/j.jviromet.2005.05.005 (2005).

55 Nie, J. et al. Quantification of SARS-CoV-2 neutralizing antibody by a pseudotyped virus-based assay. Nat Protoc 15, 3699–3715, doi:10.1038/s41596-020-0394-5 (2020).

56 Letunic, I. & Bork, P. Interactive Tree Of Life (iTOL) v5: an online tool for phylogenetic tree display and annotation. Nucleic Acids Res 49, W293–W296, doi:10.1093/nar/gkab301 (2021).

57 Pettersen, E. F. et al. UCSF ChimeraX: Structure visualization for researchers, educators, and developers. Protein Sci 30, 70–82, doi:10.1002/pro.3943 (2021).

